# Effects of MRI on an Injectable Hydrogel with Magnetically Alignable Microstructures for Oriented Cell Growth

**DOI:** 10.64898/2026.04.17.719294

**Authors:** Yas Oloumi Yazdi, Tanya Bennet, Andrew Yung, Kirsten Bale, Alex Pieters, Iryna Liubchak, Anna A. Meyer, Tara M. Caffrey, Stefan Reinsberg, Laura De Laporte, John D.W. Madden, Karen C. Cheung

## Abstract

Injectable biomaterials with aligned microstructures play a critical role in tissue engineering and drug-delivery applications where control over the position and orientation of cells and nano/micron-scale architectures enhance intervention efficacy. Patients are often subject to MRI scans; for patient safety and treatment efficacy, we investigated the effects of MRI on a biomaterial treatment consisting of aligned magnetic microstructures being developed for guiding cell growth. Under MRI exposure, potential movement of aligned structures could be detrimental to nearby cells, and potential MRI-induced heating could adversely affect traumatized tissue. In this work, the alignment state and heat conduction of such a treatment were studied using a 9.4 T preclinical MRI. The treatment comprises short magnetic rod-shaped polycaprolactone fibers (rods) with embedded magnetic nanoparticles in a surrounding hydrogel (gelatin methacrylate), with rod alignment observed before and after a 45-minute MRI scan. No change in rod alignment state was observed, and no heat generation was measured. A theoretical framework was developed which supports the experimental observation that the biomaterial is stable under MRI. This work can be extended to other biomaterial systems with aligned architectures used in tissue engineering applications such as spinal cord, muscle and tendon.

## 1. Introduction

Microstructure alignment plays a critical role in a range of biomedical applications where spatial control over cells and biomolecules is essential for promoting functional tissue organization ^1,2^. Aligned architectures have been shown to influence cell migration, orientation and differentiation, and are particularly valuable in tissue engineering strategies aimed at regenerating anisotropic tissues such as muscle ^3–5^, tendon ^6,7^, nerve ^8–10^, and myocardium ^11,12^. Approaches that enable microstructure alignment in situ, such as through external magnetic fields (MFs), are especially promising for minimally invasive therapies ^13–16^. One such approach leverages magnetically alignable microstructures (microscale rods) embedded in a hydrogel matrix called an Anisogel ^8–10^, which can be delivered in injectable form. Once injected, the gel fills the target space, and the embedded microstructures can be aligned using an externally applied MF ^17–19^. After microstructure alignment, the hydrogel is crosslinked to maintain structural orientation, which has been shown to orient cell growth. One example application is an injectable Spinal Cord Injury (SCI) treatment for which in vitro work shows promise^8–10^.

Our group is working towards a SCI treatment, comprising magnetically alignable microstructures in a hydrogel precursor solution, that would be injected into the cavity that forms at the injury site. The injected material fills the cavity and provides a medium for neuron regeneration. The microstructures contain superparamagnetic iron oxide nanoparticles (SPIONs) and can be aligned in an external MF. Once they are aligned and the surrounding precursor solutions cross-linked, the microstructures will stay in place and guide neuron regrowth across the injury site, as we have demonstrated in vitro previously ^8^.

In clinical settings, patients are subject to routine MRI scans to assess their condition, and to monitor treatment progress. These strong MFs have the potential to change rod alignment, making the approach incompatible with MRI imaging. In addition, for patient safety, it is necessary to investigate the effect of the strong Direct Current (DC) and gradient fields of MRI scans on the magnetic rods in crosslinked gel, both with regards to their aligned state and potential heat generation. Any movement of the microstructures will affect nearby cells, while heat may be detrimental to the recovery of injured and inflamed tissue. This work is the first experimental test of the behavior of a hydrogel-rod matrix described above, in an MRI scan: looking into the behavior of the magnetic microstructures in the crosslinked hydrogel both in terms of their alignment state, and potential heating. The hydrogel used in this study is gelatin methacrylate (referred to as GelMA in this work), which is crosslinked using a Ruthenium-based photo initiator. GelMA was selected for our work due to its biocompatibility, cell-adhesive properties, programmable mechanical properties and photopolymerization abilities ^20–23^; by modifying different parameters such as polymer concentration, degree of functionalization, and the photo crosslinker used (along with the photo crosslinker concentration, exposure wavelength and time) the aforementioned properties can be tuned for optimal cell growth ^22,24^. The selection of ruthenium and sodium persulfate as the photo crosslinker system allows us to crosslink the GelMA formulations using visible blue light (400-450 nm light) for improved clinical relevance as opposed to Ultraviolet light ^25,26^. The magnetic microstructures (rods) used are polycaprolactone (PCL) fibers ^17,19,27^. These PCL rods have been tested and shown to influence the growth direction of cell culture as can be seen in ^19^. The overall hydrogel-rod system concept has demonstrated oriented cell growth in our previous works presented in ^8–10^. Additional details on the samples used can be found in *Methods* (Section 4).

Following a 45-minute MRI scan designed to resemble a clinical spinal cord MRI in a 9.4 T preclinical MRI scanner (details in *Methods*, Section 4.3 and 4.4), the alignment state of the rods remained unchanged (*Results*, Section 2.1) and no heating was observed up to a scale of 0.1°C (*Results*, Section 2.2). These results supported by a developed theoretical framework (*Methods*, Section 4.1 and 4.2, and *Discussion*, Section 3). The use of a 9.4 T preclinical scanner represents the upper bound of results expected to be seen in a clinical MRI, operating at MFs ≤ 3 T ^28,29^.

## 2. Results

The results are presented in terms of the two studies conducted: Section 2.1 covering the Alignment State Study and Section 2.2 covering the Heat Generation Study. These results will be discussed in the *Discussion* and *Methods* (Sections 3.1, 4.1 for the Alignment State Study; Sections 3.2, 4.2 for the Heat Generation Study). Details on the sample preparation (Section 4.3), experimental setup (Section 4.4) along with the relevant analysis (Section 4.5) for each study can be found in *Methods* (Section 4).

For the GelMA formulation used in this work (6% GelMA, 50% DoF crosslinked using blue light with a photo initiator containing Ru and SPS), in the high MF strength (9.4 T) MRI scanner used to conduct a 45-minute scan (described in the *Methods*, Section 4.4.1), rods unaligned and aligned both parallel and perpendicular to the large Direct Current (DC) field do not experience a magnetic torque large enough to overcome the mechanical constraint of the covalently crosslinked hydrogel and cause a permanent change in their alignment (*Results*, Section 2.1; *Methods* and *Discussion* with theoretical framework Section 4.1, 3.1). In a separate study, the temperature change (*Results*, Section 2.2) due to eddy currents is measured to be less than 0.1 °C in an isolated sample (1 mL sample volume), comparable in volume to that expected for injection into a human spinal cord lesion; these results are supported by a theoretical framework presented in *Discussion* and *Methods* (Sections 3.2, 4.2).

### 2.1 Alignment State Study

Results from the visual (image overlay and Histogram of Oriented Gradients, HOG) and numerical quantification methods (Root Mean Squared Error, RMSE; Correlation Coefficient, CC) discussed in *Methods* (Section 4.5.1) can be seen in Fig. 1, for one representative well of each condition, outlined in Fig. 7(a).

**Figure 1.**
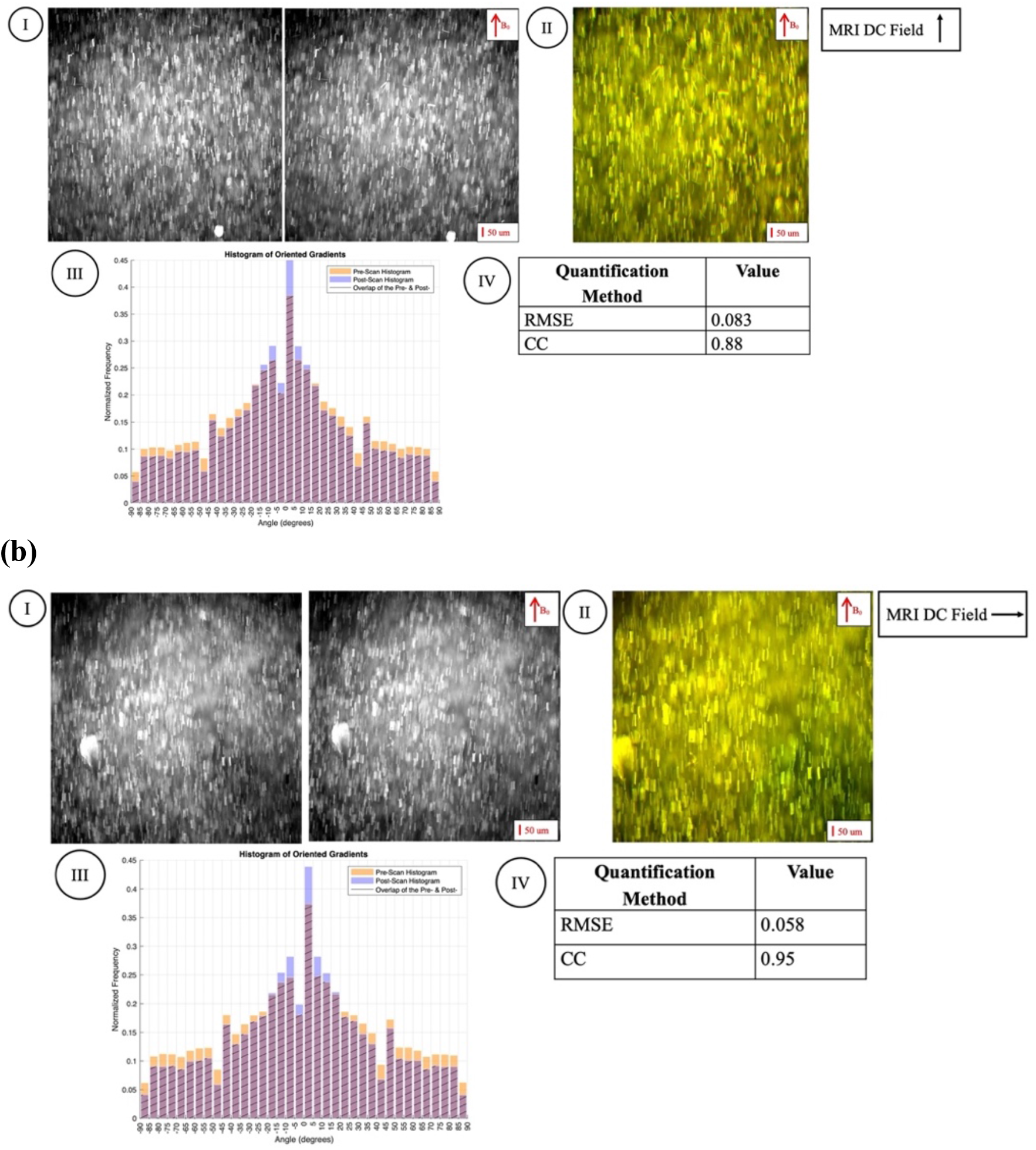
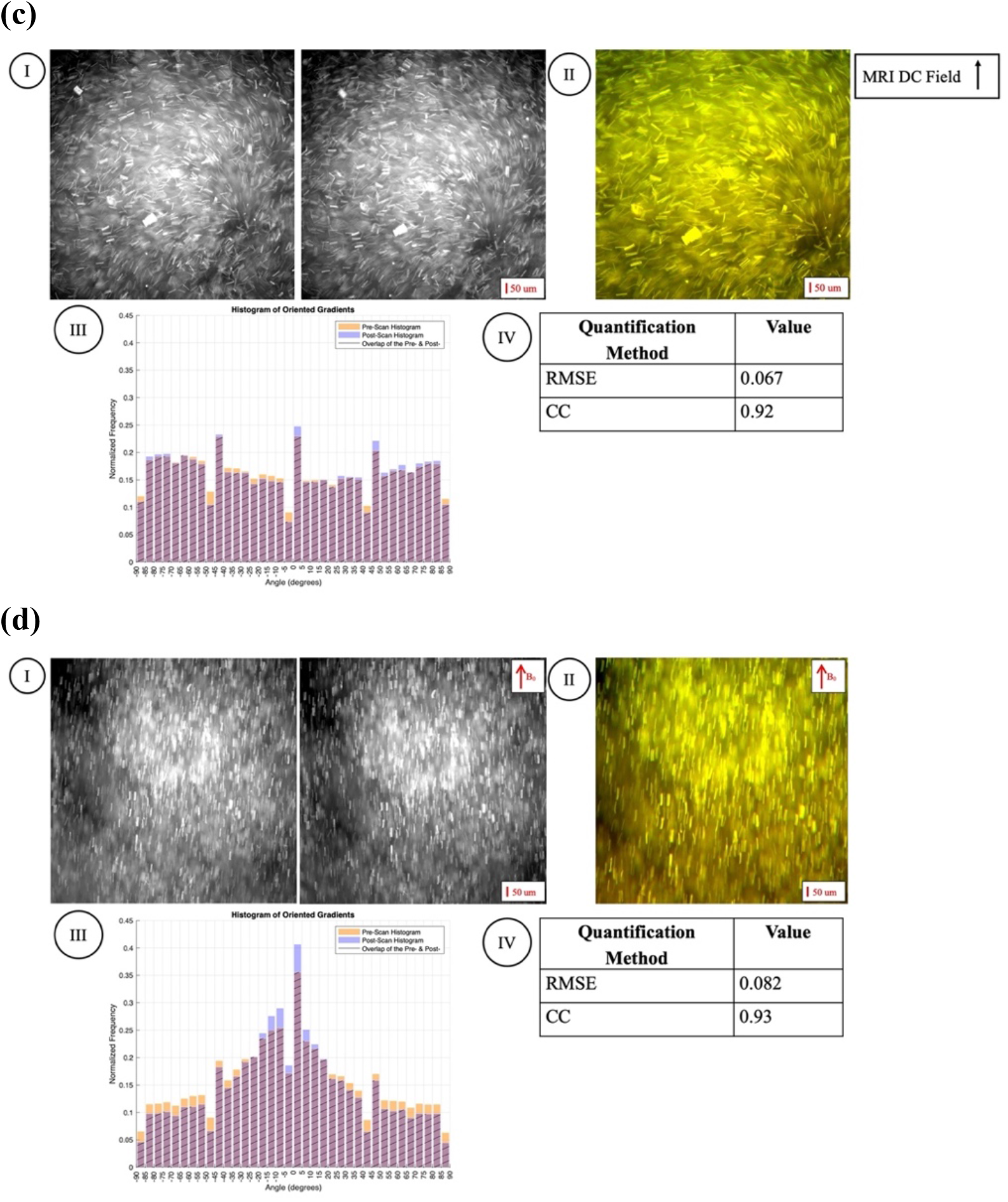
Image analysis results for representative sample wells. **(a)** Chip 2: rod aligned parallel to *B*_*0*_ of the MRI, **(b)** Chip 4: rods aligned perpendicular to *B*_*0*_ of the MRI, **(c)** Chip 1: unaligned rods **(d)** Control Chip (Chip 9): aligned rods, no MRI exposure. The four different blocks in each image are: (I) Pre (left) and post (right) MRI scan wide field fluorescence images of one of the wells of different sample conditions. *B*_*0*_ in the marked images indicates the direction of sample alignment. (II) Overlay of the pre (red) and post (green) scan images. The overall yellow color of the image indicates good overlap between the two. (III) Overlaid HOG (Histogram of Oriented Gradients) of pre (orange) and post (blue) scan images; the overlap can be seen in purple, with black hatch lines. (IV) Numerical quantification values for sample well. RMSE: Root Mean Squared Error. CC: Correlation Coefficient.

From the visual comparison metrics in Fig. 1(I, II), the before (I, left) and after images (I, right) show little change in all cases, as can be seen from the overlay of the pre- (in red) and post-(in green) scan images showing a consistent yellow color indicating good overlap between the two images. The alignment maintenance is also evident by comparing the before and after images by eye. Comparing the HOGs of the MRI-exposed samples to the control samples, we see similar patterns in the HOGs generated, indicating that the MRI does not affect the permanent alignment of the rods. As these are wide field fluorescence images captured on a Nikon Ti2 Epifluorescence microscope (the rods contain nile red as a fluorescent dye ^19^), the images acquired are not in a specific plane; they are capturing signals from the entirety of the sample thickness. The slight difference in the pre- versus post-images could therefore be due to the imaging of slightly different areas in the sample wells. Therefore, from both the HOGs and overlaid images, we can conclude that the samples were unaffected by the MRI scans despite the incredibly high field (9.4 T). Next, the numerical quantification methods will be considered (Figure 2, mean values can be found in *Supplementary Information*, Section IV).

**Figure 2.**
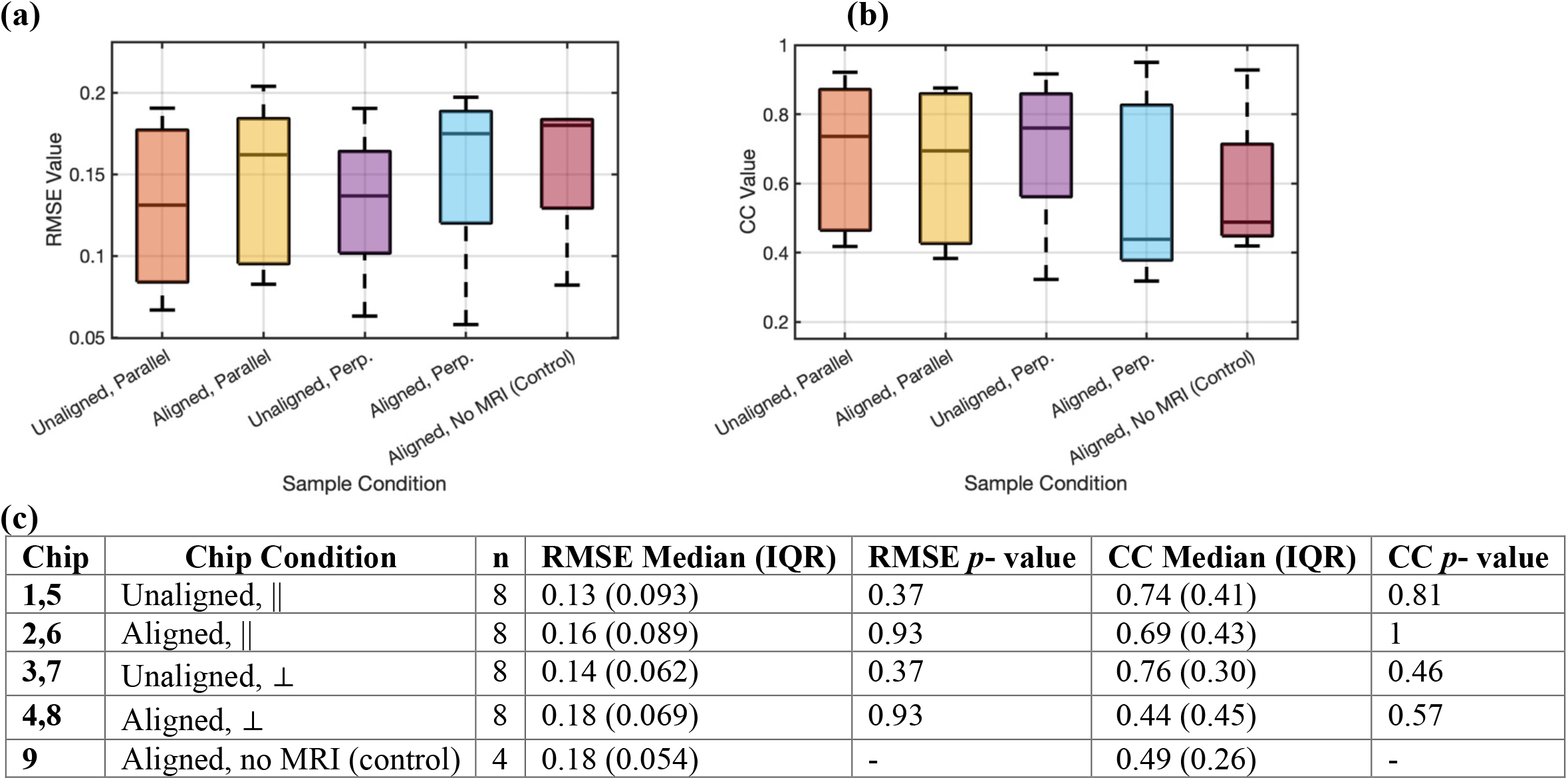
Alignment State Study Numerical Quantification Results. **(a)** Box plot of the numerical quantification value, RMSE (range: 0 to 1). **(b)** Box plot of the numerical quantification value, CC (range: -1 to 1). **(c)** Wilcoxon rank-sum test results comparing control chip (chip 9) quantification values with those of chips 1-8, which underwent the MRI scan. This data is presented along with the median, with IQR referring to the interquartile range. *p-* value <0.05 is considered as statistically significant.

From the numerical quantification values (Figure 2), the Wilcoxon rank-sum test (Mann-Whitney U test) was used to compare the samples that underwent an MRI scan with the control samples. This is a non-parametric test comparing the median of two datasets ^30^. It was used as the two samples are independent with a distribution that may not necessarily be normal. The datasets are small, making the median comparison a more effective metric compared to the mean. The results showed no statistically significant difference between the MRI-exposed samples and the control (Figure 2(c)). The overlap of the boxplots in Fig. 2(a,b) support the null results of the statistical test. Therefore, both the visual and numerical quantification methods for comparing the pre- and post-MRI scan images indicate that the samples have undergone no permanent change in their alignment due to the MRI scan.

### 2.2 Heat Generation Study

The average temperature change values across the three scans can be seen in Fig. 3; results from each individual scan can be found in the *Supplementary Information* (Section V). The results of the temperature recording for the fiber optic probe measuring ambient air temperature within the MRI RF coil can be seen in Fig. 3(a). Although the conditions were kept the same as much as was possible across the three trials, there are slight differences in the air temperatures both inside and outside the RF coils of the MRI scanner across the three trials. As can be seen from Fig. 3(b), there is some overlap between the standard deviation of the recorded measurements, indicating that the temperature values of some of the samples are not meaningfully different from the other samples. No increase in temperature was observed following 45 minutes of scanning, as will be discussed in the *Discussion* (Section 3.2) using the theoretical framework developed in *Methods* (Section 4.2).

**Figure 3.**
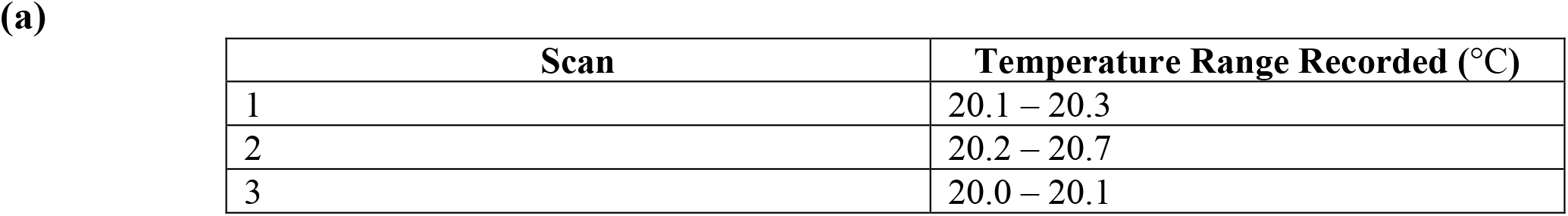
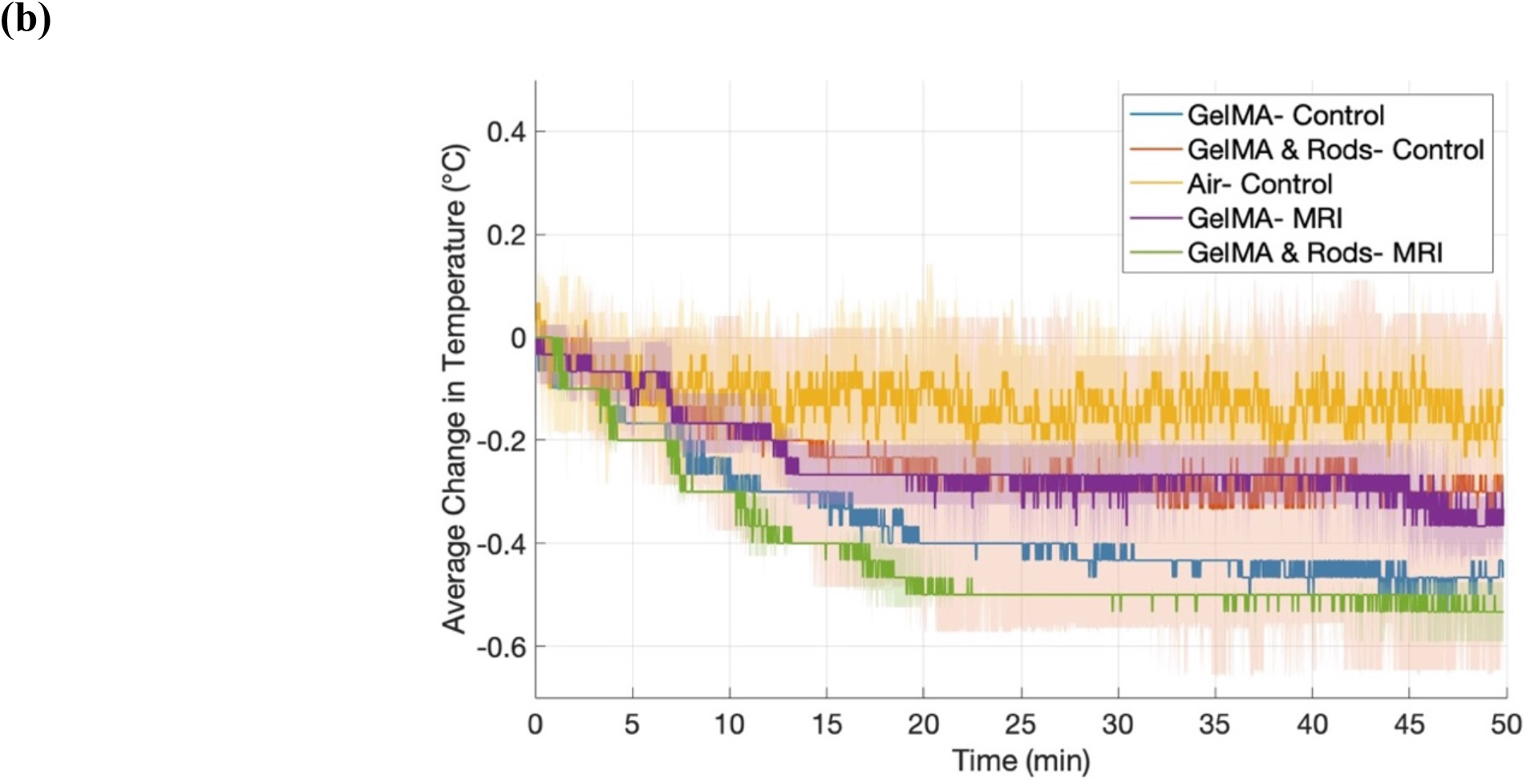
Heat Generation Study Results. **(a)** Temperature of fiber optic probe in RF coil of MRI scanner next to the experimental samples. **(b)** Average temperature change of the samples over the course of a 45-minute MRI scan. The time spans to 50 minutes as there are pauses in between scans. The sampling rate of the measurements was 1 per second. The change in temperature values used to calculate the average across the three scans are relative to the starting temperature of each sample, respectively. The shaded regions depict the standard deviation for each sample type across the three scans.

## 3. Discussion

### 3.1 Alignment State Study

As can be seen from visual comparison methods in Fig. 1 presented in the *Results* (Section 2.1), and the numerical quantification method results (Figure 2), experimentally, there were no changes in the alignment state of the rods due to the MRI scan. Using the framework presented in the *Methods* (Section 4.1), the theory behind this result for a single rod is discussed. There are two aspects to consider with regards to the movement of the rods due to the applied MF: (a) Comparing the timescale of the rapidly switching fields with the timescale needed for the rods to move in the gel, (b) Comparing the magnetic torque (*τ*_*mag*_) due to the applied MF with the elastic, viscous, and inertial torques of the gel (*τ*_*elastic*_, *τ*_*viscous*_, and *τ*_*inertia*_, respectively). Point (a) only needs to be considered for the oscillating radiofrequency (RF) and gradient fields, whereas points (a, b) are relevant for all field types of the MRI scanner. The initial framework for the discussion below is developed in the *Methods* (Section 4.1). A full list of all variables used in this work, along with their definitions and units, is provided in Supplementary Table S1.

#### 3.1.1 Rod Movement due to DC Field

Considering (b), comparing *τ*_*mag*_ due to the applied MF with *τ*_*elastic*_, *τ*_*viscous*_, *τ*_*inertia*_, this will be done for the large uniform DC field that is always ON. As presented in the *Methods* (Section 4.1), if the rods do not move with this large magnitude field, then they will not move due to the gradient and RF fields that are smaller by three orders of magnitude. Considering the torques involved ^31–36^, the manner in which the *τ*_*elastic*_, and *τ*_*viscous*_ interplay depends on the gel being used. Results from amplitude and frequency sweeps (*Supplementary Information*, Section II) are used to characterize the hydrogel formulation; details on the rheological test conducted can be found under *Gel Characterization* (Section 4.3.2). From the amplitude sweeps (Supplementary Fig. S3(a)), the gel elastic shear (storage) modulus (*G’*) in the linear viscoelastic region (LVR) is calculated as 12.06 ±7.75 Pa; the loss tangent and phase angle values (Supplementary Fig. S3(c) and (d), respectively) are below 0.45 (unitless for the loss tangent and in radians i.e. around 25° for the phase angle) thus indicating a more elastic behavior of the photo-crosslinked hydrogel ^373839^. This behavior is also observed in the frequency sweep results (Supplementary Fig. S3(b)), where the photo-crosslinked GelMA is behaving as a viscoelastic solid material ^40^; these results allow us to model the gel using the Kelvin-Voight model ^32,34,40^. Using the frequency sweep, the zero-shear viscosity (*η*_0_) is calculated as 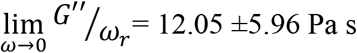, where G’’ is the gel loss modulus and *ω*_*r*_ is the angular frequency (using the *G’’* at *ω*_*r*_= 0.1 rad/s) ^34^. As the photo-crosslinked GelMA behaves as a viscoelastic solid; this allows us to use the Kelvin-Voight model with a spring and damper in parallel to describe the hydrogel ^323440^. Within this framework, the elastic behavior of the gel is presented via the elastic spring and the viscous behavior through the damper, as such the differential equation governing the movement of a single rod can be written as ^31–35^:

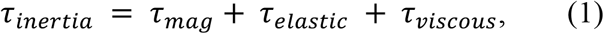

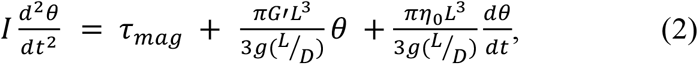

where *L* is the rod length, D is rod diameter, g(L/D) is a dimensionless function of the anisotropy ratio (L/D = *p*), *η*_0_ is the gel zero-shear viscosity, I is the rod moment of inertia, and *θ* is the rod angle of deflection with respect to its initial angle (*β*_0_); *β*_0_ is relative to the applied MF direction. This model is depicted in Fig. 5.

*τ*_*mag*_ depends on the rod saturation magnetization field (in fact of the SPIONs). In this study, the MRI applies a 9.4 T DC MF. Using the Langevin Magnetization curve provided by the SPION manufacturers, the SPIONs in this work saturate at ∼ 0.6 T, with a saturation magnetization of ∼ 50 -70 emu/g; for the calculations here, the average value of 60 emu/g is used (EMG 1200, Ferrotec; Source: Ferrotec Datasheet). For the rods, the saturation field is 0.6*ϕ*_*SPION*_ where *ϕ*_*SPION*_ is the SPION volume fraction in the rod ^41 42^. This value is smaller than the DC MRI field as the volume fraction will always be ≤1; the rods will be in their saturation regime in the scanner. *τ*_*mag*_ is therefore defined as equation (3), with the unit conversation for the saturation magnetization in equation (4). Details on the equations can be found in *Methods* (Section 4.1).

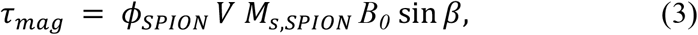

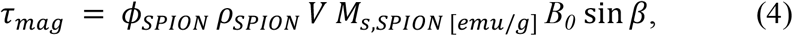

where *ρ*_*SPION*_ is the SPION density and *M*_*s,SPION* [*emu*/*g*]_ is the saturation magnetization of the SPIONs [emu/g], *B*_0_ is the applied MF, *V* is rod volume, and *β* is angle between the applied MF and the direction of magnetization (ellipsoid axis of symmetry). SPION density and volume fraction calculations can be found in the *Supplementary Information*, Section VI. If the applied field is less than the rod saturation magnetization field, the rod demagnetizing factors and magnetic susceptibility need to be calculated (*Methods*, Section 4.1; *Supplementary Information*, Section VI). From the torque equations presented, the two angles (*θ* and *β*) are linked using *β* = *β*_0_– *θ*.

For the *τ*_*elastic*_ and *τ*_*viscous*_ term, *G’* and *η*_/_were determined above; *g(L/D)* from equation (2) is calculated using equation (5) as the anisotropy ratio *p* = L/D = 10 (between 2 and 20 for the rods) ^31,43^:

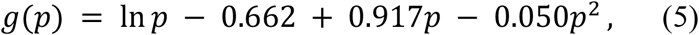

Considering *τ*_*inertia*_, as the system described is at static equilibrium, the angular acceleration of the rods goes to zero. Thus, the inertial torque term is zero, and the differential equation becomes a first order differential; rewriting it as a function of the rod retardation angle, *β* (equation (6)):

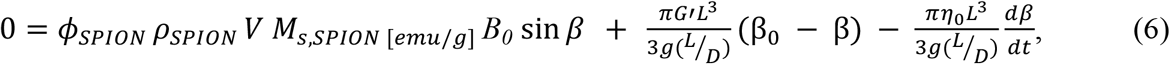

The differential equation is solved (MATLAB), and the results replicating the 9.4 T field can be seen in Fig. 4(a). As can be seen in Fig. 4(a), when rods are aligned parallel and perpendicular to the DC field, the deflection angle is 0° as was observed experimentally. The rods experience maximum torque at 45° initial angle relative to the applied MF. This results in a maximum deflection angle of 0.51°. Rods with initial angles between 0-45° and 45-90° experience a smaller deflection. This simulated maximum deflection value is a very small amount (<1°) at an applied field of 9.4 T which is not used clinically. Clinical scanners typically operate at 1.5 to 3 T ^28,29^; the differential equation can be solved for the upper bound of 3 T. The results for the 3 T applied MF can be seen in Fig. 4(b) with the largest deflection angle being 0.16° for a starting rod angle of *β*_0_ = 45°. For a 1.5 T applied field, the largest deflection angle is 0.082°. Therefore, it can be concluded that for the hydrogel and rod combination tested, it is expected that there will be minimal observed deflection of the rods due to the DC MRI MF, based on the rheological and inertial properties of the gel and rods. For other gel and rod combinations, rheological tests need to be conducted to determine the nature of the hydrogel; rod properties will also play a factor in determining any potential changes in alignment due to an MRI scan. For instance, assuming the same rods are used, any changes in the mechanical properties of the gel (*G’* and *η*_0_) given it has the same viscoelastic nature will require rerunning the simulation and potential further experiments depending on the deflections seen in the simulated results. In this hypothetical case, for the same *η*_0_, a decrease in *G’* by a factor of 2 (*G’* ∼ 6 Pa) in a 9.4 T MRI scanner will result in a maximum deflection angle of 1.02° for *β*_0_ = 45°. A gel with such *G’* is characterized as extremely soft; therefore, for such a case, additional MRI experiments would need to be conducted to determine if any rotation is observed. In cases where the simulated deflection is less than 1.00 ° i.e. in stiffer gels (*G’* > 10 Pa), any observed deflection would likely be minimal as was observed experimentally in this work. Lastly, if administered as a treatment for oriented cell growth in the case of spinal cord injury, most of the rods are expected to be aligned along the direction of the applied DC MRI field and will experience no deflection. Meanwhile, if the rods need to be aligned in a different direction for applications other than spinal cord injury treatment, the maximum deflection experienced if they are held in place by the gel would not be very substantial due to clinical MRI (at 3 T DC field: <0.16°). Additionally, these results are based on assumptions and approximations that have been made throughout this work and may not precisely characterize the hydrogel-rod environment. As such, it’s important to acknowledge that the Kelvin-Voight differential equation-based model used in this work is a simplified model of viscoelasticity and thus may not fully capture complex hydrogel behaviors. Other complex models of the gels such as the standard linear solid model (presented in ^44,45^) which serve as more realistic models of the biomaterial’s creep and relaxation can be considered for future work as needed.

**Figure 4.**
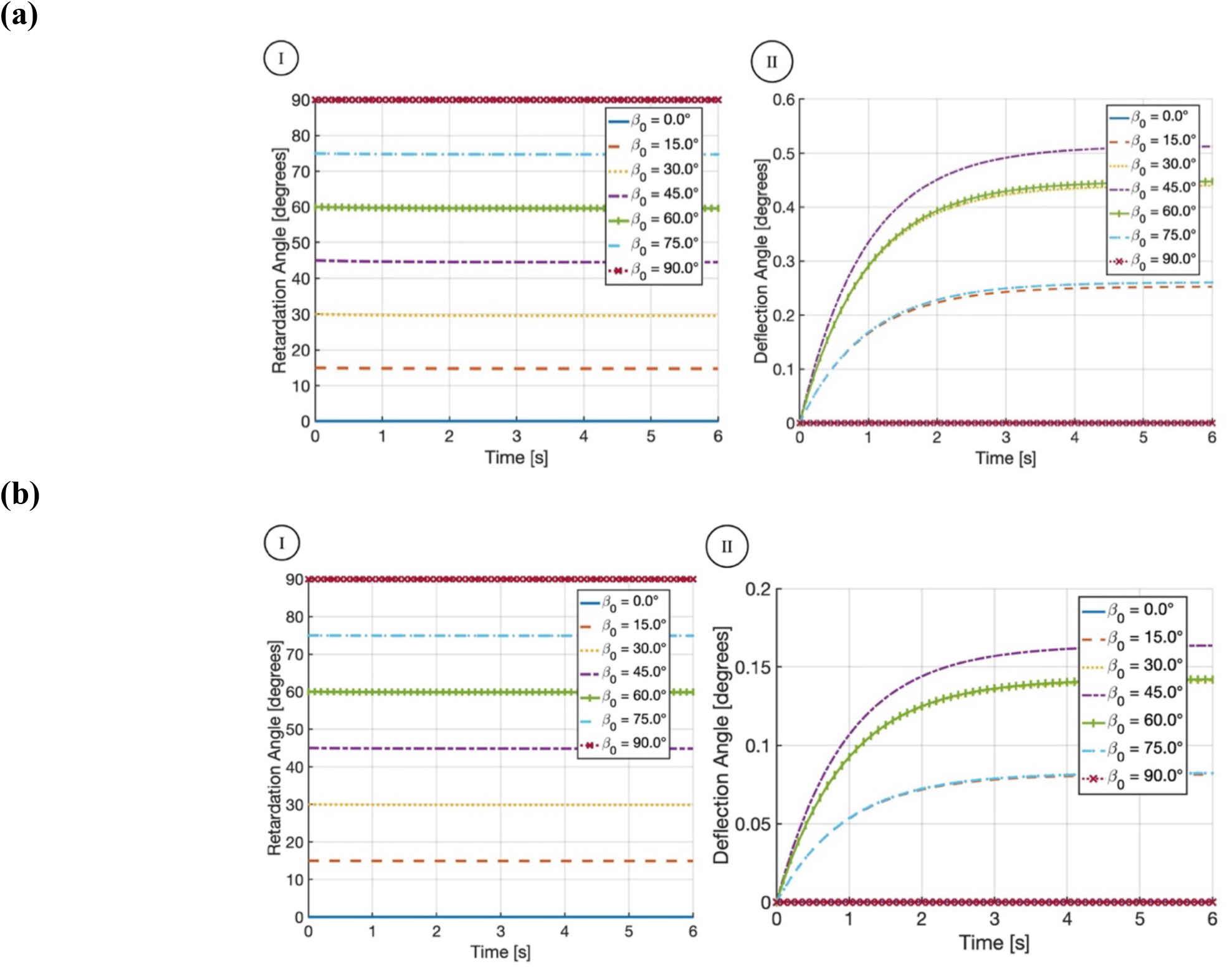
Simulation Results. Rod rotation due to **(a)** 9.4 T, **(b)** 3 T applied MFs at different initial angles (*β*_0_); this initial angle is with respect to the applied MF direction. For each field, (I) Rod retardation angle (*β*) with respect to the static applied MF; (II) rod deflection angle with respect to the rod initial angle.

#### 3.1.2 Rod Movement due to RF and Gradient Fields

As presented in the *Methods* (Section 4.4.1), the timescale of the gradient fields is ∼100 µs, with a maximum magnitude of 667.94 mT/m (2.9 mT in the experimental sample sizes used). Meanwhile, for the RF fields, the timescale (defined in terms of pulse width, *PW*) had a range of 4.3 µs ≤ PW ≤ 3.5 ms, with magnitude ∼µT scale (<= 0.48 mT). The repetition time between the pulse patterns (*TR*) had a range of 4.2 ms – 2.5 s. Thus, although the exact timescale for rod movement is not known, as they are being held in a covalently crosslinked matrix of the gel, it is not expected that the rod movement time is shorter than the micro- to milli-second scale pulses applied by the gradient and RF fields. Additionally, from the simulated results presented in Fig. 4, the time for rod movement to occur in the crosslinked hydrogel is in the scale of seconds, further reiterating the point that frequency of oscillations will not lead to significant rod movement and/or rotation. As such, it can be concluded that the gradient and RF pulses are not expected to affect the rod alignment state, as was observed experimentally. The theoretical framework underlying rod movement supports the experimental results: the PCL rods in the specific GelMA formulation used do not experience a change in their alignment state due to the MFs of a 45-minute MRI scan conducted using a 9.4T preclinical MRI scanner. Changes in the surrounding hydrogel rheological properties will affect the timescale of rod movement; the MATLAB simulation developed needs to be modified accordingly to determine the timescale within which the rods respond to the applied MRI MFs.

### 3.2 Heat Generation Study

From Fig. 3 in the *Results* (Section 2.2), the samples are not experiencing a rise in their temperature due to the MRI RF field. In fact, the samples appear to be dropping in temperature. This was observed in the control and the experimental samples. MRI scanners are cooled through liquid helium ^46^. While monitoring the temperature of components within the scanner (including the six gradient coils and two chiller waters in the scanner) during the scans, values ranging from 13.4 °C to 25.7 °C were noted, with the other components having temperature values within this range. Thus, although the RF fields may be causing heating of the samples, the cooling observed due to the scanner environment occurs at a larger scale compared to any potential heating.

The maximum expected temperature change of the samples can be estimated using equation (7) derived in the *Methods* (Section 4.2) ^36,47^:

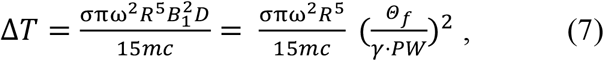

where *σ* is the electrical conductivity of the substance in the sphere, *D* is the duty cycle of the RF pulse, *ω* is the angular frequency of the RF MF, *B*_1_ is the RF MF amplitude, *R* is the radius of the sphere that approximates the shape of the sample, *m* is the mass of the sample, *c* is the specific heat capacity of the sample, 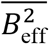 is the weighted sum of the RF pulses in a single MRI scan, *B*_*eff*_ is the effective rectangular pulse amplitude, *Θ*_*f*_ is the flip angle (radians), *PW* is the pulse width and *γ* is the gyromagnetic ratio (Hz/T) ^47^. The flip angle and pulse width are different for each pulse type in the different scans conducted (*Supplementary Information*, Section VII). *ω* of the scanner was 400 MHz. The cylindrical samples were approximated as spherical samples: original dimensions of radius 5.64 mm and height 10 mm, estimated as sphere of radius 5.64 mm. Regarding *c* and *σ*: as the GelMA is only at 6%, it is quite diluted and has a large DMEM volume (Dulbecco’s Modified Eagle Medium, high glucose, HEPES, no phenol red, Gibco, ThermoFisher Scientific). DMEM is primarily composed of water. As such, *c* of water is used in the calculations (4184 J/ (kg K)). *σ* of DMEM is measured to be 1.005 S/m (Electrical Conductivity Meter: Hach HQ430D) at 19.4°C. The total temperature rise for each scan can be seen in *Supplementary Information*, Section VII (Table S4). For the entire 45-minute scan, the expected temperature rise calculated theoretically is 1.33 x10^-3^ °C. Thus, the theoretical results support the experimental results that the heating of the samples is < 0.1°C, which is the measurement resolution of the temperature probes used, and the scale of the cooling observed in the experimental samples. These results are for a 9.4 T scanner with a higher field and SAR compared to clinal MRI scanners, thus representing an upper bound of the heating expected in a clinical MRI scan.

Lastly, *σ* of surrounding spinal cord tissue can be compared with that of DMEM to compare expected temperature rise. *σ* of human cerebrospinal fluid (CSF) is 1.45 S/m at room temperature (25°C) and 1.79 S/m at body temperature (37°C) ^48^. Other sections of the spinal cord have the following *σ*: grey matter 0.333 S/m, white matter 0.143 S/m, dura 0.030 S/m, vertebrae 0.006 S/m, disks 0.200 S/m measured at a body temperature not under 36 °C, less than 24 h post-mortem ^49^. As can be seen, some regions of the spinal cord have a higher, and some a lower *σ* compared to DMEM. As such, during the MRI scan, higher *σ* regions will experience a greater temperature rise, while those with a lower *σ* will experience a lower temperature rise compared to the administered hydrogel combination treatment. Additionally, in this experimental setup, the samples used were surrounded by PDMS, which is thermally insulative. If the treatment were to be administered to a patient, then the sample volume would be surrounded by human tissue which itself heats up in an MRI scan ^50^.

### 3.3 Conclusion

Through this study, the effects of an MRI scan on a matrix of crosslinked hydrogel with magnetic microstructures was studied. The effect was investigated with regards to both the alignment state as well as heating of the hydrogel-rod system. The alignment state of the rods remained unchanged during a 45-minute scan designed to resemble a clinical spinal cord MRI conducted in a 9.4 T preclinical MRI scanner. No heating was observed up to a scale of 0.1°C. A 9.4 T scanner has a higher field and SAR compared to clinal MRI scanners that operate at MFs ≤ 3 T. As such, the results of this study represent an upper bound of magnetic forces on the rods and potential heating of the system that are expected in a clinical MRI scan.

Future studies include conducting the alignment studies with additives such as growth factors, or anti-inflammatory drugs added to the hydrogel. Addition of these is aimed to enhance cell growth in the hydrogel. These additives however may affect the mechanical properties of the gel, as well as its crosslinking efficacy. More complex hydrogel models may be required to effectively capture intricate behaviors of the gel. These drugs may also affect the electrical conductivity of the sample due to the presence of additional ions, which may result in heating above the acceptable safe range. Both the alignment state and heating behavior can first be assessed using the respective theoretical frameworks developed in this work, before a final treatment combination can be assessed using an MRI scan. This framework supports efficient pre-evaluation of candidate treatment combinations, reducing reliance on extensive MRI-based testing that would otherwise be required to assess all possible configurations. Lastly, with regards to the heat studies, additional studies can be conducted with the samples placed in MRI phantoms with conductivity similar to human tissue resulting in a more physiologically representative system ^36,51^. In this work, a baseline has been developed for studying the behavior of a hydrogel-magnetic microstructure combination in an MRI scan. Such approaches are highly applicable in various fields, and this study will serve as a baseline for other work aiming to assess their respective systems with regards to their MRI-safety to ensure patient welfare.

## 4. Methods

### 4.1 Alignment State Study Framework: How can the magnetic fields of an MRI scan affect the alignment state of magnetic microstructures?

A patient subject to an MRI scan is exposed to three different fields: the main DC magnetic field (MF, *B*_*0*_), a radiofrequency field (RF, *B*_*1*_), and gradient fields (*G*_*x*_, *G*_*y*_, *G*_*z*_) ^52^. *B*_*0*_ is parallel to the scanner bore’s axis and results in the net magnetization of the tissue and any implanted materials. The RF coils induce a secondary MF perpendicular to *B*_*0*_, which allows for the creation and detection of a transverse magnetization; they are the main source of potential heating in the scan. The gradient coils are used for spatial encoding of the signals and switch on and off with varying amplitudes and periods ^52^. When considering the effect of the MFs on the rod alignment state, the three MFs must be considered: *B*_*0*_, gradient fields (*G*_*x*_, *G*_*y*_, *G*_*z*_), and *B*_*1*_. The effect each has depends on the initial alignment state of the rods. In this work, the initial alignment state of the rods is presented with respect to the applied MF direction; the behavior of the rods due to the applied MFs and the mechanical constraints of the gel will be considered in terms of a torque balance as is presented below.

The DC field, *B*_*0*_, is uniform and always present. To determine whether or not the rods will move due to *B*_*0*_, the magnetic torque (*τ*_*mag*_) of the rods due to the applied MF must be compared with the elastic restoring torque (*τ*_*elastic*_), viscous (*τ*_*viscous*_), as well as inertial (*τ*_*inertia*_) torques of the gel. The magnetic torque component is the cross product of magnetization (**M**) of the rod with the applied MF (**B**_**0**_). The magnetization can be written as (equation 8) ^41^:

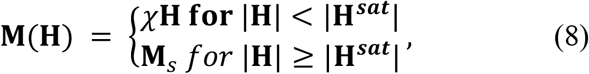

where **H** is the total internal MF, ***H***^**sat**^ is the internal MF at saturation, and **M**_*s*_ is saturation magnetization of the rods. The magnetic torque ranges from 0 to a maximum value for initial rod angles of 0° to 45°, respectively. This torque is generated due to magnetic anisotropy within the rods; it has been observed experimentally that the rods under a spatially uniform external MF will rotate to align their long (major) axis with the direction of the applied MF ^19,41^.

Approximating the rods as ellipsoids, the magnetic torque on the rods in the linear regime prior to magnetic saturation (|***H***| < |***H***^**sat**^|) can be expressed using equation (8) ^36,41^:

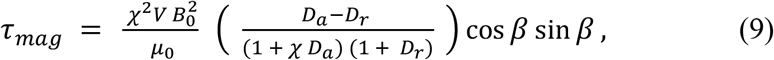

where χ is the rod magnetic susceptibility, *V* is the rod volume, *B*_*o*_ is the applied MF, *μ*_/_ is the vacuum magnetic permeability, *β* is angle between the applied MF and the direction of magnetization (ellipsoid axis of symmetry), *D*_*a*_ is the demagnetizing factor along the axis of symmetry, and *D*_*r*_ is the radial demagnetizing factor ^53^. After the saturation magnetization has been reached (|***H***| ≥ |***H***^**sat**^|), the magnetic torque can be defined using equation (10):

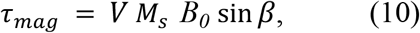

where in this case the assumption has been made that the magnetization direction of the rod is along its long axis. This assumption simplifies further calculations and serves as an upper bound for the torque experienced by the rods due to the induced magnetization from the applied MRI applied MF.

Considering the *τ*_*elastic*_, *τ*_*viscous*_, and *τ*_*inertia*_, the exact combination within which they affect rod movement depends on the rheological properties of the surrounding crosslinked hydrogel; the gel used in this study will be characterized in *Gel Characterization* (Section 2.1.2). *τ*_*elastic*_, *τ*_*viscous*_, and *τ*_*inertia*_ can be defined by equations (11-13) ^31–35^:

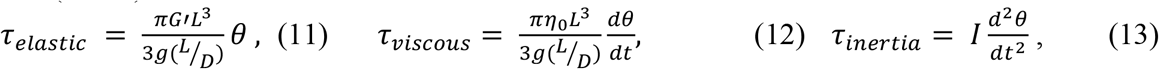

where *G’* is the gel elastic shear (storage) modulus, *L* is the rod length, D is rod diameter, g(L/D) is a dimensionless function of the anisotropy ratio (L/D = *p*), *η*_0_ is the gel zero-shear viscosity, I is the rod moment of inertia, and *θ* is the rod angle of deflection with respect to its initial angle. This initial angle, *β*_0_, is relative to the applied MF direction. From the equations for torque presented above, the two angles are linked using *β* = *β*_0_– *θ*. Additional details for this specific study were presented earlier in the *Discussion* (Section 3.1).

Looking at *G*_*x*_, *G*_*y*_, *G*_*z*_, *B*_*1*_, there are two aspects to consider: (a) Comparing the timescale of the rapidly switching fields with the timescale needed for the rods to move in the gel; (b) comparing the magnetic torque of the rods due to the applied MF with the mechanical constraint of the gel in terms of the restoring torques as mentioned above. With regards to (a), the timescales will be discussed briefly in the *Discussion* (Section 3.1), but they are different depending on the hydrogel-rod and MRI system used. Considering (b), if the rods are not expected to overcome the mechanical barrier of the gel even when the largest magnetic torque is applied due to the large DC field mentioned above, then the gradient and RF fields are too small in magnitude to be able to cause movement of the rods. Two main assumptions will be made in this work with regards to the rods moving forward: the rods are treated as rigid structures that do not bend, with a concentration that is sufficiently dilute that there is no interference either magnetic or mechanical amongst them. As such, calculations for a single rod will extend to all the other rods.

We have now outlined the framework regarding the alignment state of the rods. A full list of all variables used in this work, along with their definitions and units, is provided in Supplementary Table S1. Figure 5 depicts the torques acting on a rod in a crosslinked gel: the torque due to the applied MFs (*τ*_*mag*_) of the MRI scanner, as well as the restoring torque of the gel in the elastic regime (*τ*_*elastic*_), the viscous torque (*τ*_*viscous*_) that arises due to the viscous behavior of the gel, as well as the inertial torque (*τ*_*inertia*_). The mechanical magnetic torque of the rod (*τ*_*mag*_) due to the applied MRI MFs will be balanced by the torques from the gel-rod interactions and the rod inertia (*τ*_*elastic*_, *τ*_*viscous*_, *τ*_*inertia*_). When balance is achieved with minimal angular deflection, *θ*, the rods remain stable and do not change in orientation. In addition, the elastic torque will return the rods to their original orientations, once the field is removed. If rod-gel combinations are used where the gel has significant creep (in general if it is behaving as a liquid), then we may see reorientation, particularly over long periods, since response is not dominated by elasticity. If rods are shorter than the hydrogel cross-link spacing (typically nanometer scale) or pore size ^54,55^ then re-orientation may be observed even if the hydrogel on average is elastic. Rod-gel interfacial interactions (interfacial adhesion and surface chemistry of the rods) may also play a role in the relative freedom to move ^56–58^.

**Figure 5.**
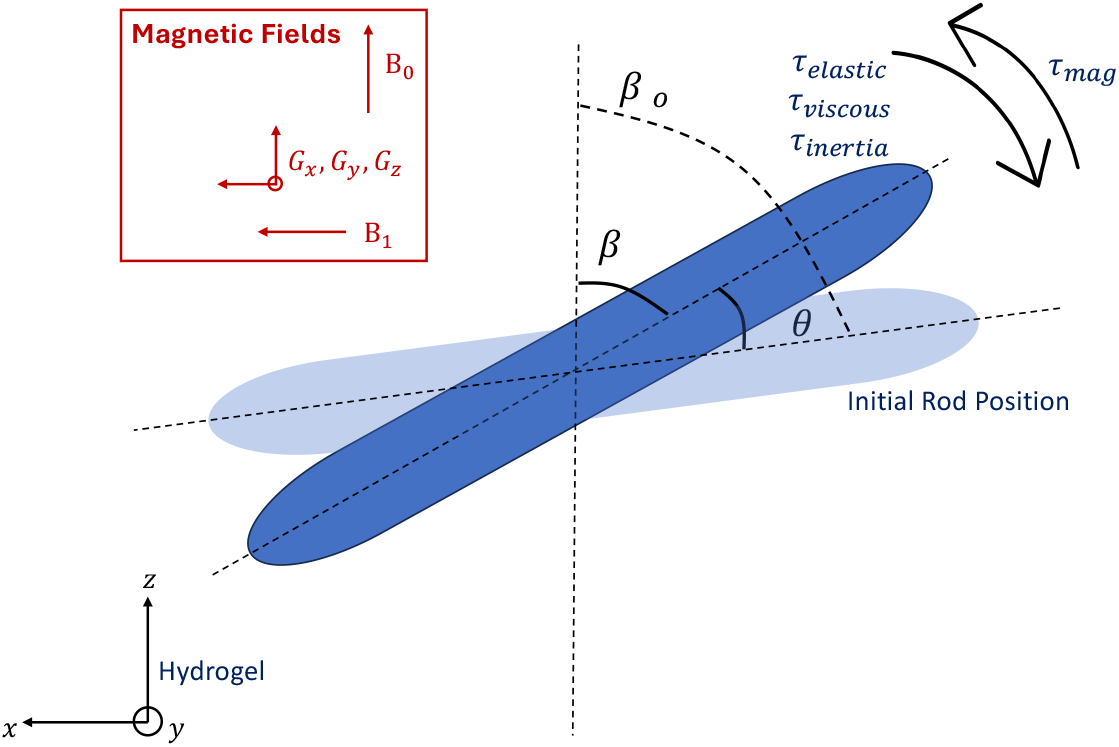
Diagram depicting the torques acting on a single rod in a gel when placed in an MRI scanner, where *β*_0_ is the initial rod angle with respect to the applied MF direction, in this case looking at the effect of *B*_*0*_ on rod alignment state, *β* is the current rod angle with respect to the applied MF direction, and *θ* is the rod deflection angle with respect to its initial position. As such, we can define *β* = *β*_0_– *θ*.

### 4.2 Heat Generation Study Framework: What may cause heating of samples during an MRI scan?

The RF field of an MRI scan can cause heating of conductive samples due to Joule heating. The changing MF of the RF waves causes an electric field in conductive samples by Faraday’s Law. This induced electric field results in the flow of charges, resulting in a closed loop of current called eddy current. The eddy current in turn causes joule heating ^36,59^. The power, *P*, generated in a spherical sample of radius, *R*, due to the RF field induced eddy current can be estimated from the expression (equation 14):

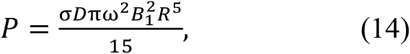

where *σ* is the electrical conductivity of the substance in the sphere, *D* is the RF pulse duty cycle, which is the rate of the RF pulse duration, *ω* is the angular frequency of the RF MF, and *B*_1_ is the RF MF amplitude. While the sample is not in general spherical (as will be presented in the *Methods* Section 4.1, the samples used are short, nearly equidimensional cylinders), the equation establishes the magnitude of the effect. From the power, the upper bound of temperature rise can be calculated using the time for each scan (assuming no power is lost, and all is absorbed as heat), by equation (15):

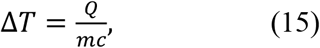

where *Q* is the amount of heat energy absorbed by the sample, *m* is the mass of the sample, and *c* is the specific heat capacity of the sample.

Each protocol in an MRI scan has different RF pulse types in each repetition time, TR (time from one set of RF pulses to the next). To calculate the total power for each scan, the different duty cycles (*D*) and RF MF amplitudes (*B*_1_) of each pulse type (excitation pulse, refocusing pulse, and fat suppression pulse) are needed. This is the 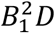 term; for a single scan type, this is equivalent to the weighted sum of the *D* and 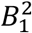 for each pulse type. This can be achieved using equations (16,17) ^47^:

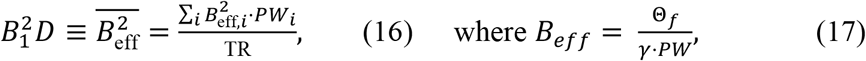

and where 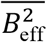 is the weighted sum of the RF pulses in a single MRI scan, *B*_*eff*_ is the effective rectangular pulse amplitude, *Θ*_*f*_ is the flip angle (radians), and *γ* in the gyromagnetic ratio (Hz/T). The flip angle (*Θ*_*f*_) is the angle of rotation of the net magnetization of the sample relative to the DC MF (B_o_) due to RF excitation pulse ^47^. Putting this all together (equation (18)):

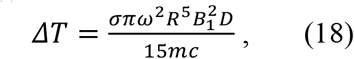

The details of this calculation specific to MRI scans in this study were outlined in the *Discussion* (Section 3.2).

### 4.3 Sample Preparation

The hydrogel used in this study is 50% degree of functionalization (DoF) gelatin methacrylate (PhotoGel50%DS, CELLINK; referred to as GelMA in this work), which is crosslinked (using blue light) with a photo initiator containing tris(2,2’-bipyridyl) ruthenium (II) (Ru) and sodium persulfate (SPS) (Sigma-Aldrich). The steps for hydrogel sample preparation are as follows: (a) Prepare 15% w/v stock of GelMA. This is done by adding 3.35 mL of phosphate buffered saline (PBS) to 500 mg of freeze-dried GelMA. This sample is then magnetically stirred at 200 rpm at 45°C. It is protected from ambient light in this time (e.g. by wrapping in aluminum foil) to prevent premature crosslinking. (b) Prepare 10 mg/mLRu stock in PBS. (c) Prepare a fresh 10 mg/mL SPS stock. This stock solution is made fresh for each experimental run. Once the stock solutions of GelMA, Ru, and SPS are prepared, a solution of 6% w/v GelMA with 0.2 mM Ru/2.0 mM SPS is prepared. The magnetic microstructures (rods) used are polycaprolactone (PCL) fibers ^17,27^. These rod-shaped fibers (50 µm length, 5 µm diameter) have a magnetic response due to the incorporation of SPIONs. The SPIONs allow for the alignment of rods using milli-Tesla (mT) scale externally applied MFs. Details on PCL rod fabrication are briefly discussed under *PCL Rod Fabrication* (Section 2.1.1) ^8–10,18,27^. Based on previous work exploring the relationship between the rod concentration inside the gel and aligned neuronal outgrowth, rods are added to achieve 1% v/v concentration ^60,61^. In the case of the Alignment State studies, the sample is then pipetted in a PDMS (Polydimethylsiloxane) 4-well chip (Figure 6(a)). Each chip contains four wells of 20 *μ*L volume within which the sample is injected into. The wells are then covered with PBS to prevent the samples from drying out. In the case of the Heat studies, these samples are injected into PDMS human-sized mock spinal lesions (MSLs) using a 1 mL syringe (22G dispensing tip); these PDMS MSLs (Figure 6(b)) have an enclosed 1 mL “cavity”. Details on the fabrication of the PDMS 4-well chip and MSL can be found in the *Supplementary Information* (Section I). Once injected, the PDMS structure (either MSL, or 4-well chip) with the GelMA with/without rods combination is placed under a light source (initially OFF) and a MF source. First the sample is aligned for 60 seconds by the field, before the light source is turned ON for another 60 seconds. If the GelMA does not contain any rods, it is not subjected to the MF. The MF source used to achieve alignment is composed of two stacks of two 38.1 mm diameter, 19.05 mm thick N52 permanent neodymium magnets (K&J Magnetics Inc.) placed in a parallel configuration. The magnets are raised 20 mm from the base, with a 7.2 cm magnet centre-to-centre distance. This setup generates a MF strength of 42 mT at the centre of the magnetic configuration. An LED light source is used for photo-crosslinking the GelMA (PLS-0455-010-C, MIGHTEX Precision LED Spotlight, *λ*= 450 nm, 63.7 mW/cm^2^ power density). Sample preparation steps can be seen in Fig. 6(c) ^8–10,18,27^.

**Figure 6.**
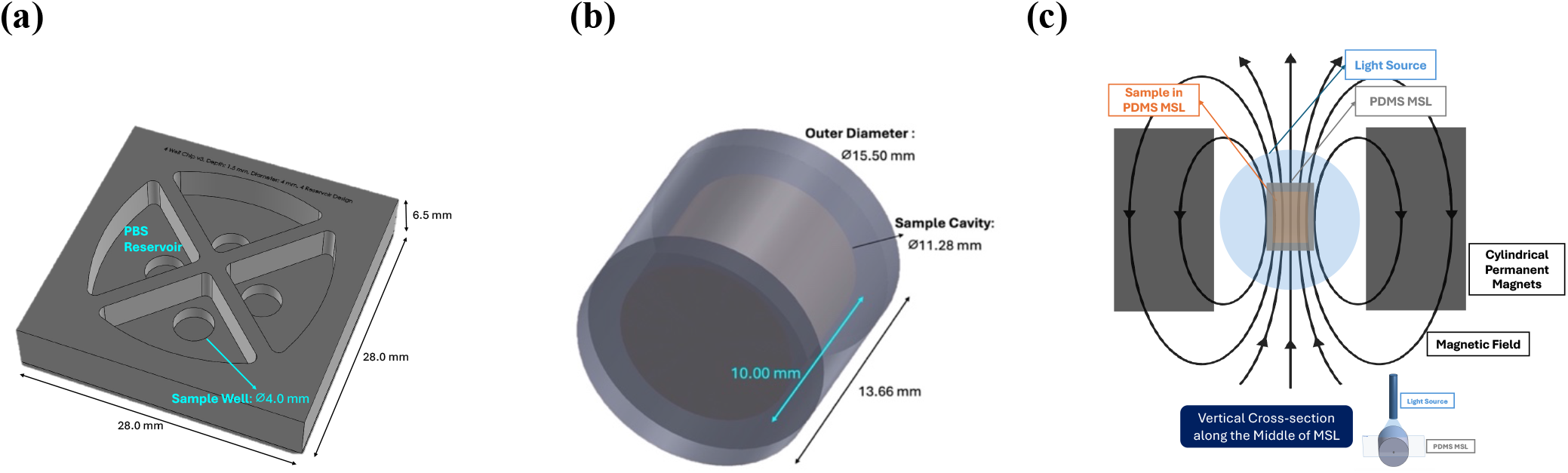
Sample Preparation Summary. **(a)** Alignment State study setup: Drawing of 4-well PDMS chip design. **(b)** Heat generation study setup: Drawing of Mock Spinal Lesion (MSL) design. **(c)** MSL under light and MF source. Similar setup is used for the 4-well PDMS chip. Images not drawn to scale.

#### 4.3.1 PCL Rod Fabrication

The short magnetic PCL rods were produced by wet-dry fiber spinning. For spinning solution preparation, chloroform, SPIONs (1.70 mg/mL, equals 0.99 weight (w/w) % of solid polymer content) and nile red (0.01 mg/mL) were mixed by vortexing; 17 wt/vol% PCL was then dissolved in the premixed solution overnight on a roller mixer. The spinning solution was transferred into a glass syringe (TLL1002, Hamilton) equipped with a custom-bent U-shaped, blunt cannula (Sterican 21G, 120 mm, B. Braun) and connected to a syringe pump (AL-1000, World Precision Instruments). The pump was positioned behind an ethanol bath such that the U-shaped cannula was immersed in the bath. For spinning, the extruded filament was picked up with a tweezer at the cannula tip, slowly drawn upwards and placed onto a rotating aluminum drum (custom-built setup with drum diameter: 5 cm), on which the filament was collected in an aligned manner. Before spinning, the drum was wrapped in aluminum foil and coated with a dried layer of optimal cutting temperature (OCT) compound. Spinning was conducted with a flow rate of 0.3 mL/hr and drum speed of 300 rpm. When fiber collection was finished, the spun fibers were coated with another layer of OCT and dried overnight. The OCT-embedded aligned PCL fibers were then harvested from the drum and sectioned into 50 μm thick slices using a cryostat microtome (Leica CM1950, equipped with FEATHER C35 microtome blades). Harvested sections were collected in a tube and purified by centrifugation, followed by four washing steps with water to remove the OCT. The fiber dispersions were then disinfected by UV irradiation under continuous stirring for 3 hrs ^19^.

#### 4.3.2 Gel Characterization

From the torque equations presented in the *Discussion* and *Methods* (Section 3.1 and 4.1), the shear elastic (storage) modulus (*G’*) and zero-shear viscosity (*η*_/_) of the gel must be determined for use in the Alignment Study theoretical framework developed. Additionally, to determine the mechanical model of viscoelasticity to be used for the photo-crosslinked hydrogel, the loss tangent and phase angle (tan *δ*= G’’/G’ where *G’’* is the loss modulus, *δ* is the phase angle, and G’’/G’ is the loss tangent) which serve as indicators of how elastic or viscous a material is behaving also need to be determined. The storage modulus (*G’*) is measured within the linear viscoelastic region (LVR) of the gel via an oscillatory strain sweep (0.1-10% at 5 rad/s; n=4) ^62,63^. The viscosity as well as the loss tangent of the gel are determined using a frequency sweep (0.1-10 rad/s at 0.5% strain; n = 7). The GelMA samples are prepared 24 hours prior to the rheological characterization. They are stored in PBS at 37 °C for this duration to allow them to reach equilibrium swollen state. The crosslinked hydrogels are then transferred to a DHR30 (Discovery Hybrid Rheometer 30, TA Instruments) for characterization. The frequency sweep data was taken on 2 separate days (day 1: 3 samples; day 2: 4 samples). The amplitude sweep data was taken on the same ‘day 2’ samples (n =4). On Day 2, the order of the sweeps was first a non-destructive frequency sweep followed by the amplitude sweep to confirm that the strain selected in the frequency sweep falls within LVR.

### 4.4 Experimental Setup

#### 4.4.1 Alignment State Study Experiments

The studies in this work were conducted using a 9.4 T preclinical scanner (Bruker BioSpec 94/30) at the Centre for Comparative Medicine (CCM, UBC). Clinical scanners typically operate at 1.5 or 3 T ^28,29^. Using a preclinical scanner allowed us to study the upper bound scenario, where the MF strengths, maximum gradient slew rate and amplitude, and SAR (Specific Absorption Rate) exceed the current clinical scenario. The MRI scan lasted a total of 45 minutes, as is typical in a clinical spinal cord MRI ^64^. We used a diverse scan protocol to explore a range of imaging tasks that may be seen on a clinical scanner. The scans conducted were designed by the UBC MRI Research Centre to mimic clinical spinal cord scans, and are as follows:

1. Initial three-slice localizer (FLASH). This is used to identify the sample at the isocenter of the scanner; 43 s duration.
2. Three “fast spin echo” (RARE) scout images, one for each orthogonal direction; 1 min 20 s, 1 min 20 s, and 40 s duration.
3. Two isotropic-resolution gradient echo (FLASH) scans; 1.5 min, 6 min duration.
4. Fast echo-planar imaging (EPI) scan, typical for fMRI experiments. This scan has strong gradients switching rapidly allowing for rapid acquisition; 7.5 min
5. Diffusion tensor echo-planar imaging (DT-EPI). This scan type shares the same strong and fast gradient pulses as EPI, but also includes strong gradient pulses played out less frequently to sensitize signals to the diffusivity of water; 14 min
6. Ultrashort echo time (UTE) in both 2D and 3D. This scan has a very short echo time for direct imaging of tissues with very short T2 (transverse) relaxation times, such as bone and certain areas of the brain, particularly myelin; 1 min duration.

The sample conditions tested are shown in Fig. 7(a); these were prepared as was outlined in the *Sample Preparation* section: 6% w/v GelMA with 0.2 mM Ru/2.0 mM SPS and 1% v/v concentration PCL magnetic rods. Both aligned and unaligned rod samples were imaged. Samples aligned and parallel to *B*_*0*_ represented the expected scenario within which a SCI patient with the potential treatment would be subject to, i.e. the rods are aligned along the SC. Unaligned samples represented the scenario within which alignment was not achieved in the patient. Samples aligned perpendicular to *B*_*0*_, i.e. all rods aligned perpendicular to the intended tissue guidance direction, were also included in the study as an additional reference for ensuring no rod movement is experimentally observed when the model and theory predict no movement (either translation and/or rotation). Each condition had two repeat chips (total of 8 chips per MRI scan), with each chip containing four identical wells, i.e. there is a total of 8 samples per condition. A control chip (chip 9) that contained aligned rods but was not subject to an MRI scan was also used; therefore, there are 4 control samples. The samples were prepared on Day 0 and imaged under a Nikon Ti2 Epifluorescence microscope. The prepared samples were then stored in a 37°C incubator overnight. On Day 1, the samples were transported to CCM to be placed in the MRI scanner and then taken back to the lab and imaged under the microscope (10X magnification). The layout of the PDMS chips can be seen in Fig. 7(b).

**Figure 7.**
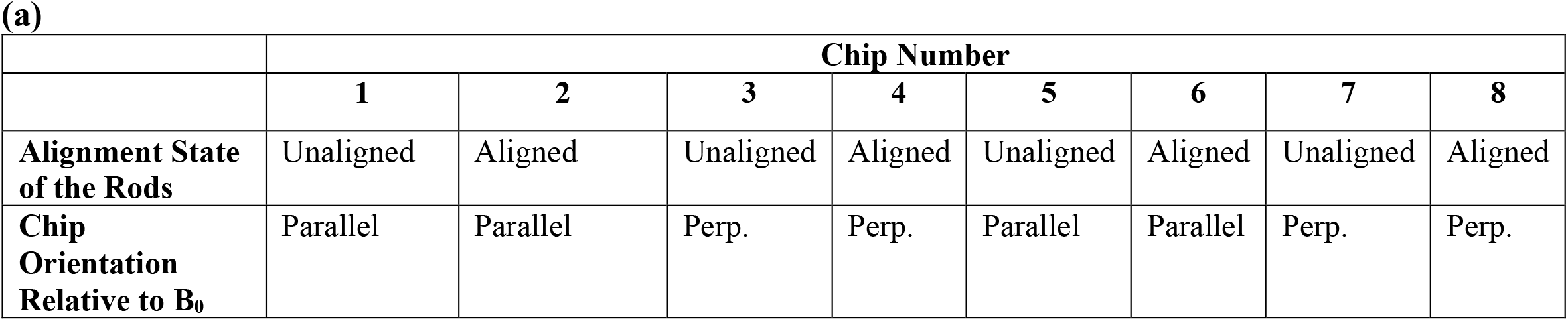
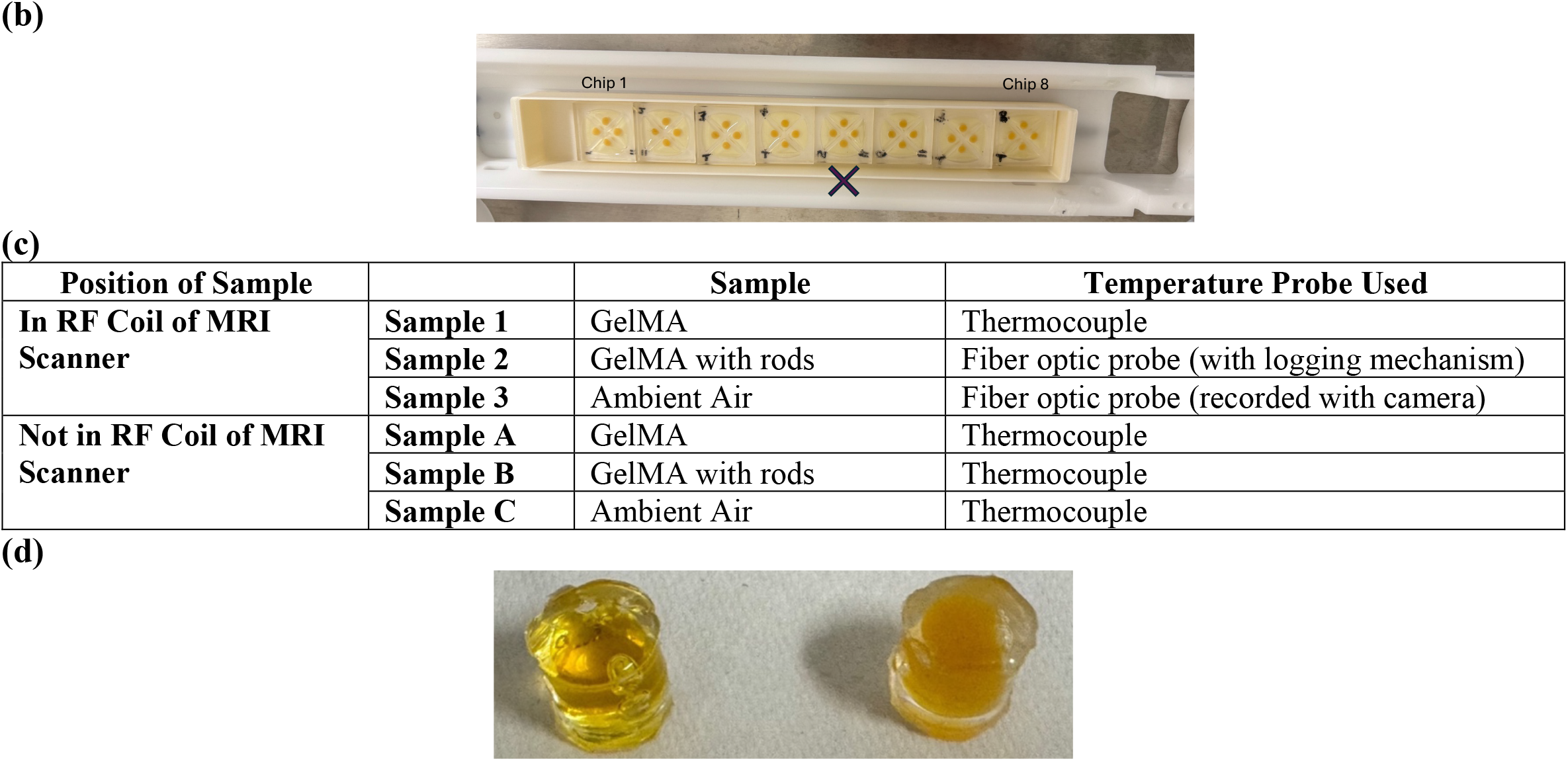
Sample Setup. **(a)** Conditions tested in the Alignment State studies. Chips 5-8 are repeats of chips 1-4. Aligned samples were placed parallel and perpendicular to the large DC field (B_0_) of the MRI scanner; the perpendicular samples were achieved by rotating the PDMS chips by 90°. In the case of the unaligned samples, the “orientation” refers to the PDMS chips being rotated as was done for the aligned samples to ensure consistency in sample preparation. Perp.: perpendicular. **(b)** Layout of chips within MRI scanner. The MRI isocenter is marked with an **x** (chip 5). **(c)** Samples tested in the MRI scan to assess generated heat. Within each scan, the temperature of three samples within the RF coil of the MRI scanner, and three samples outside the RF coil but within the magnet bore were monitored. Three scans, each with freshly prepared samples, were conducted on the same day **(d)** MSL fabricated out of PDMS. The left is filled with GelMA, and the right is filled with GelMA and rods. The temperature probes were inserted into small holes on the curved surface of the MSLs.

#### 4.4.2 Heat Generation Study Experiments

Heat studies were conducted using the same 9.4 T preclinical scanner (Bruker BioSpec 94/30) at CCM. The same scans as the Alignment State studies were conducted, as they were designed to mimic clinical MRI spinal cord scans by the UBC MRI Research Centre. The sample conditions tested are shown in Fig. 7(c); these were prepared as was outlined in the *Sample Preparation* Section above: 6% w/v GelMA, 0.2 mM Ru/2.0 mM SPS with/without 1% v/v concentration PCL magnetic rods. The PDMS MSLs with the injected and crosslinked samples can be seen in Fig. 7(d). Each scan was limited to three samples due to the availability of MRI-compatible temperature probes: one thermocouple (RTP-102-B, Instruments for Small Animals I4SA) and two fiber optic probes (FTP-SA3-ST1, I4SA). One fiber optic probe lacked a logging mechanism; its readings were recorded manually from the display using a camera and subsequently transcribed. The same control samples were used; these control samples were placed within the bore of the MRI scanner but not subject to the RF and gradient fields as they were placed outside the RF coil. As MRI scanners are cooled by liquid helium, these control samples would be subject to the same potential cooling, but not the heating sources. The temperature of the samples not in the scanner was measured using a different thermocouple sensor probe (Ultra-Fine, TI-SP-K, 2-9249-01, MISUMI) and datalogger (GRAPHTEC GL2000). Three identical scans were conducted. The samples were prepared fresh for each scan: the samples were placed in the scanner after at least 10 minutes post-preparation before waiting another 10 minutes before the MRI scan was started to ensure temperature stabilization.

### 4.5 Analysis Performed

#### 4.5.1 Alignment State Study Experimental Analysis

As mentioned in the *Methods: Experimental Setup* (Section 4.4.1), pre- and post-scan images are obtained on a Nikon Ti2 Epifluorescence microscope (10X magnification). These images are acquired through a Section in the middle of the sample which is in focus. As sample height and volume is not changing pre- and post-MRI scan, a Section at a similar location in the gel is being imaged. Following area-based image registration using pixel-intensity ^65^, several methods are used: two visual methods, and two numerical quantification methods, as can be seen from Fig. 8(a). An image registration step involving rotation and translation was applied to facilitate direct comparison between pre- and post-MRI images. This step does not impact the validity of the rod alignment analysis. In the case of image rotation, detection of changes in rod orientation would only be affected if all rods experienced identical rotation due to the MF and the image registration applied an equivalent counterrotation. With respect to translation, if rod displacement had occurred during the MRI exposure, it would have resulted in a mismatch between corresponding structural features in the pre- and post-images; this mismatch was not observed. Thus, the registration procedure allows image comparison without introducing artefacts that would confound the analysis of rod alignment. The visual methods include the overlay of the registered images, as well as the generation and overlay of a Histogram of Oriented Gradients (HOG, MATLAB) ^66^. HOG is typically used for feature extraction, but in this work, it is used to determine the angle of the gradients allowing us to observe the angle of the rods in images; summary of the steps can be seen in Fig. 8(b). To quantify differences between pre- and post-MRI images, two complementary methods were employed: Root Mean Squared Error (RMSE) and Correlation Coefficient (CC) in 2D. Each method emphasizes different image features, enabling a more comprehensive and unbiased assessment. RMSE provides a pixel-wise comparison based on absolute intensity differences, reflecting the deviations across the entire image. As a traditional “error-sensitivity” metric, RMSE is sensitive to shifts in pixel intensity, but it does not account for structural or perceptual similarity between images. CC is also a pixel-based comparison tool quantifying the linear correlation (relationship) between the pixel-intensities of two images ^67^; it is widely used in template matching and image registration ^68,69^. The value generated gives a linear indication of the similarity between the two images ^70,71^. This multi-metric approach improves the robustness and interpretability of image comparisons, particularly in detecting subtle or spatially complex changes in rod alignment. A summary of the two methods can be found in the *Supplementary Information* (Section III). The results of these comparison methods, both visual and numerical, are shown in the *Results* (Section 2).

**Figure 8.**
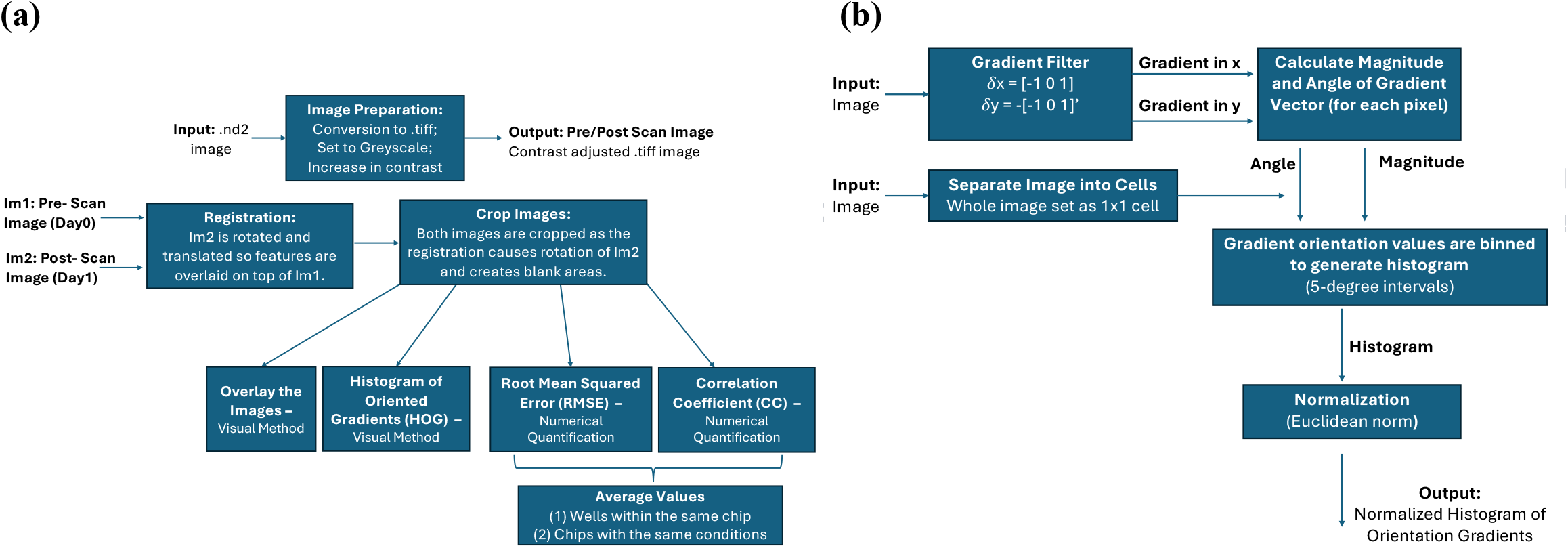
Analysis Steps Summary. **(a)** Flowchart depicting the analysis steps conducted. **(b)** Flowchart depicting HOG generation steps.

#### 4.5.2 Heat Generation Study Experimental Analysis

The temperatures of the samples are plotted over time (MATLAB). As we are interested in the change in temperature to consider the possibility of heating, the temperature change of each sample relative to its own starting temperature is plotted and that data is analyzed, as was presented in the *Results* (Section 2) and *Supplementary Information* (Section V).

## Acknowledgments

The authors are grateful to work and live on the traditional, ancestral and unceded territory of the Coast Salish Peoples, including the Musqueam and Squamish First Nations. The authors would like to acknowledge the support of the Government of Canada’s New Frontiers in Research Fund (NFRF) [NFRFT-2020-00238] as well as fellowship support through the Natural Sciences and Engineering Research Council of Canada’s (NSERC) Canada Graduate Scholarships—Master’s Program, the Li Tze Fong Memorial Fellowship and UBC Four Year Fellowship awards. This work was also supported through funding of the Senatsausschuss Wettbewerb (SAW) Leibniz-Transfer project µTISSUEfab and Leibniz Health Technologies Short Term Scientific Mission (STSM) funding. Lastly, the authors would also like to thank the UBC Soil and Biometeorology Teaching Lab (Lewis Fausak) for use of their equipment.

## Author Contributions

Y. O. Y. led the work presented in this paper, from experimental planning to data analysis; she wrote the main manuscript text and prepared most of the figures. T.B. prepared the experimental samples and prepared some of the figures in the main text and Supplementary Information. A.Y. and K.B. assisted with experimental planning as well as handling of MRI scans; along with Dr. S. R., they also developed the MRI protocols used, as well as assisted with MRI theory. A.P. characterized the gel used in the study through the rheological tests presented, with the gel formulation used being developed by I. L. The magnetic microstructures used were developed and fabricated by A. A.M. in the lab of Dr. L.D.L, who also provided manuscript feedback. Dr. T.M.C. reviewed the manuscript. The work was supervised by Drs. J.D.W. M. and K.C.C., both of whom provided contributions to experimental planning and gel design along with revising the manuscript.

## Data Availability Statement

The datasets generated during and/or analyzed during the current study are available from the corresponding author on reasonable request.

## Competing Interests Statement

The author(s) declare no competing interests.

## Notes

### Competing Interest Statement

The authors have declared no competing interest.

## References

1. Lu, K. et al. Biofabrication of aligned structures that guide cell orientation and applications in tissue engineering. Bio-des. Manuf. (2021) doi:10.1007/s42242-020-00104-5.

2. Xing, J., Liu, N., Xu, N., Chen, W. & Xing, D. Engineering Complex Anisotropic Scaffolds beyond Simply Uniaxial Alignment for Tissue Engineering. Adv. Funct. Mater. 32, (2022).

3. Volpi, M., Paradiso, A., Costantini, M. & Święszkowski, W. Hydrogel-Based Fiber Biofabrication Techniques for Skeletal Muscle Tissue Engineering. ACS Biomater. Sci. Eng. 8, 379–405 (2022).

4. Jana, S., Levengood, S. K. L. & Zhang, M. Anisotropic Materials for Skeletal-Muscle-Tissue Engineering. Adv. Mater. 28, 10588–10612 (2016).

5. Blake, C. et al. Replace and repair: Biomimetic bioprinting for effective muscle engineering. APL Bioengineering 5, 031502 (2021).

6. No, Y. J., Castilho, M., Ramaswamy, Y. & Zreiqat, H. Role of biomaterials and controlled architecture on tendon/ligament repair and regeneration. Adv. Mater. 32, e1904511 (2020).

7. Ghazvahanian, E., Shadman-Manesh, V., Sahranavard, M. & Ghorbani, F. Bioinspired and biomimetic materials/architecture in tendon and ligament regeneration. in Principles of bioinspired and biomimetic regenerative medicine (eds. Ghorbani, F., Ghalandari, B. & Liu, C.) vol. 3 621–655 (Springer Nature Switzerland, 2025).

8. Omidinia-Anarkoli, A. et al. An Injectable Hybrid Hydrogel with Oriented Short Fibers Induces Unidirectional Growth of Functional Nerve Cells. Small 13, (2017).

9. Braunmiller, D. L. et al. Pre-programmed rod-shaped microgels to create multi-directional anisogels for 3D tissue engineering. Adv. Funct. Mater. 2202430 (2022) doi:10.1002/adfm.202202430.

10. Vedaraman, S. et al. Anisometric microstructures to determine minimal critical physical cues required for neurite alignment. Adv. Healthc. Mater. 10, e2100874 (2021).

11. Kristen, M. et al. Fiber scaffold patterning for mending hearts: 3D organization bringing the next step. Adv. Healthc. Mater. 9, e1900775 (2020).

12. Baghersad, S., Sathish Kumar, A., Kipper, M. J., Popat, K. & Wang, Z. Recent Advances in Tissue-Engineered Cardiac Scaffolds—The Progress and Gap in Mimicking Native Myocardium Mechanical Behaviors. JFB 14, 269 (2023).

13. Li, K., Xu, J., Li, P. & Fan, Y. A review of magnetic ordered materials in biomedical field: Constructions, applications and prospects. Composites Part B: Engineering 228, 109401 (2022).

14. Rivera-Rodriguez, A. & Rinaldi-Ramos, C. M. Emerging Biomedical Applications Based on the Response of Magnetic Nanoparticles to Time-Varying Magnetic Fields. Annu. Rev. Chem. Biomol. Eng. 12, 163–185 (2021).

15. Wychowaniec, J. K. & Brougham, D. F. Emerging Magnetic Fabrication Technologies Provide Controllable Hierarchically-Structured Biomaterials and Stimulus Response for Biomedical Applications. Adv Sci (Weinh) 9, e2202278 (2022).

16. Beachley, V., Katsanevakis, E., Zhang, N. & Wen, X. Highly aligned polymer nanofiber structures: fabrication and applications in tissue engineering. in Biomedical applications of polymeric nanofibers (eds. Jayakumar, R. & Nair, S.) vol. 246 171–212 (Springer Berlin Heidelberg, 2012).

17. Licht, C. et al. Synthetic 3D PEG-Anisogel Tailored with Fibronectin Fragments Induce Aligned Nerve Extension. Biomacromolecules 20, 4075–4087 (2019).

18. Rose, J. C. et al. How much physical guidance is needed to orient growing axons in 3D hydrogels? Adv. Healthc. Mater. 9, e2000886 (2020).

19. Meyer, A. A. et al. Fabrication of Short Polymeric µFibers as Building Blocks for Anisotropic High-Throughput Compatible 3D Tissue Models. BioRxiv (2025) doi:10.64898/2025.12.09.692980.

20. Pepelanova, I., Kruppa, K., Scheper, T. & Lavrentieva, A. Gelatin-Methacryloyl (GelMA) Hydrogels with Defined Degree of Functionalization as a Versatile Toolkit for 3D Cell Culture and Extrusion Bioprinting. Bioengineering (Basel) 5, (2018).

21. Liang, J., Dijkstra, P. J., Poot, A. A. & Grijpma, D. W. Hybrid Hydrogels Based on Methacrylate-Functionalized Gelatin (GelMA) and Synthetic Polymers. Biomedical Materials & Devices (2022) doi:10.1007/s44174-022-00023-2.

22. Piao, Y. et al. Biomedical applications of gelatin methacryloyl hydrogels. Engineered Regeneration 2, 47–56 (2021).

23. Bennet, T. J. et al. Engineering a Multilayer Microfluidic Airway-On-A-Chip with Tunable GelMA Hydrogel for Physiologically Relevant Aerosol Exposure Studies. BioRxiv (2025) doi:10.1101/2025.11.06.687056.

24. Nichol, J. W. et al. Cell-laden microengineered gelatin methacrylate hydrogels. Biomaterials 31, 5536– 5544 (2010).

25. Elkhoury, K., Zuazola, J. & Vijayavenkataraman, S. Bioprinting the future using light: A review on photocrosslinking reactions, photoreactive groups, and photoinitiators. SLAS Technol. 28, 142–151 (2023).

26. Lim, K. S. et al. Visible Light Cross-Linking of Gelatin Hydrogels Offers an Enhanced Cell Microenvironment with Improved Light Penetration Depth. Macromol. Biosci. 19, e1900098 (2019).

27. Omidinia-Anarkoli, A. et al. Solvent-Induced Nanotopographies of Single Microfibers Regulate Cell Mechanotransduction. ACS Appl. Mater. Interfaces 11, 7671–7685 (2019).

28. Canadian medical imaging inventory 2022–2023: MRI: CMII report. (Canadian Agency for Drugs and Technologies in Health, 2024).

29. Committee on the Current Status and Future Direction of High-Magnetic-Field Science in the United States, Phase II et al. The Current Status and Future Direction of High-Magnetic-Field Science and Technology in the United States. (National Academies Press, 2024). doi:10.17226/27830.

30. Siegel, S. Nonparametric Statistics. The American Statistician 11, 13–19 (1957).

31. Chevry, L., Sampathkumar, N. K., Cebers, A. & Berret, J. F. Magnetic wire-based sensors for the microrheology of complex fluids. Phys. Rev. E Stat. Nonlin. Soft Matter Phys. 88, 062306 (2013).

32. Berret, J. F. Local viscoelasticity of living cells measured by rotational magnetic spectroscopy. Nat. Commun. 7, 10134 (2016).

33. Gu, Y. & Kornev, K. G. Ferromagnetic nanorods in applications to control of the in-plane anisotropy of composite films and for in situ characterization of the film rheology. Adv. Funct. Mater. 26, 3796–3808 (2016).

34. Larson, R. G. The structure and rheology of complex fluids. (1999).

35. Frka-Petesic, B. et al. Dynamics of paramagnetic nanostructured rods under rotating field. J. Magn. Magn. Mater. 323, 1309–1313 (2011).

36. Magnetic resonance procedures: health effects and safety. (CRC Press, 2000). doi:10.1201/9781420041569.

37. Nolle, A. W. Methods for Measuring Dynamic Mechanical Properties of Rubber-Like Materials. J. Appl. Phys. 19, 753–774 (1948).

38. Fuss, F. K. The Loss Tangent of Visco-Elastic Models. in Nonlinear approaches in engineering applications: applied mechanics, vibration control, and numerical analysis (eds. Dai, L. & Jazar, R.N.) 137–157 (Springer International Publishing, 2015). doi:10.1007/978-3-319-09462-5_6.

39. Seoudi, B. M., Kulik, V. M., Boiko, A. V., Chun, H. H. & Lee, I. New approach to the computation of the form factor of viscoelastic cylinders. Mechanics of Materials 41, 495–505 (2009).

40. A Basic Introduction to Rheology. (2016).

41. Kim, Y. & Zhao, X. Magnetic soft materials and robots. Chem. Rev. 122, 5317–5364 (2022).

42. Reinelt, M. et al. Simulation and experimental validation of magnetic nanoparticle accumulation in a bloodstream mimicking flow system. J. Magn. Magn. Mater. 582, 170984 (2023).

43. Tirado, M. M., Martínez, C. L. & de la Torre, J.G. Comparison of theories for the translational and rotational diffusion coefficients of rod-like macromolecules. Application to short DNA fragments. J. Chem. Phys. 81, 2047–2052 (1984).

44. Lin, C.-Y. Rethinking and researching the physical meaning of the standard linear solid model in viscoelasticity. Mechanics of Advanced Materials and Structures 1–16 (2023) doi:10.1080/15376494.2022.2156638.

45. Lakes, P. R. Viscoelastic Materials. (Cambridge University Press, 2010).

46. Huettel, S. A., Song, A. W. & McCarthy, G. MRI Scanners. in Functional magnetic resonance imaging 27–48 (2014).

47. Wang, J., Mao, W., Qiu, M., Smith, M. B. & Constable, R. T. Factors influencing flip angle mapping in MRI: RF pulse shape, slice-select gradients, off-resonance excitation, and B0 inhomogeneities. Magn. Reson. Med. 56, 463–468 (2006).

48. Baumann, S. B., Wozny, D. R., Kelly, S. K. & Meno, F. M. The electrical conductivity of human cerebrospinal fluid at body temperature. IEEE Trans Biomed Eng 44, 220–223 (1997).

49. Fernandes, S. R., Salvador, R., Wenger, C., de Carvalho, M. & Miranda, P. C. Transcutaneous spinal direct current stimulation of the lumbar and sacral spinal cord: a modelling study. J. Neural Eng. 15, 036008 (2018).

50. Bottomley, P. A. Turning up the heat on MRI. J. Am. Coll. Radiol. 5, 853–855 (2008).

51. Shellock, F. G. & Kanal, Emanuel. Magnetic resonance: Bioeffects, safety, and patient management. (1996).

52. Tourais, J., Coletti, C. & Weingärtner, S. Brief introduction to MRI physics. in Magnetic Resonance Image Reconstruction - Theory, Methods, and Applications vol. 7 3–36 (Elsevier, 2022).

53. Osborn, J. A. Demagnetizing factors of the general ellipsoid. Phys. Rev. 67, 351–357 (1945).

54. Talaei, A. et al. Optimizing the composition of gelatin methacryloyl and hyaluronic acid methacryloyl hydrogels to maximize mechanical and transport properties using response surface methodology. J. Biomed. Mater. Res. Part B Appl. Biomater. 111, 526–537 (2023).

55. Shie, M.-Y. et al. Effects of Gelatin Methacrylate Bio-ink Concentration on Mechano-Physical Properties and Human Dermal Fibroblast Behavior. Polymers (Basel) 12, (2020).

56. Wu, G. M. & Shyng, Y. T. Surface modification and interfacial adhesion of rigid rod PBO fibre by methanesulfonic acid treatment. Composites Part A: Applied Science and Manufacturing 35, 1291–1300 (2004).

57. Chhetri, S. & Bougherara, H. A comprehensive review on surface modification of UHMWPE fiber and interfacial properties. Composites Part A: Applied Science and Manufacturing 140, 106146 (2021).

58. Drzal, L. T. The role of the fiber-matrix interphase on composite properties. Vacuum 41, 1615–1618 (1990).

59. Tang, M. & Yamamoto, T. Progress in understanding radiofrequency heating and burn injuries for safer MR imaging. Magn. Reson. Med. Sci. 22, 7–25 (2023).

60. Babu, S. et al. How do the local physical, biochemical, and mechanical properties of an injectable synthetic anisotropic hydrogel affect oriented nerve growth? Adv. Funct. Mater. 2202468 (2022) doi:10.1002/adfm.202202468.

61. Rose, J. C. et al. Nerve Cells Decide to Orient inside an Injectable Hydrogel with Minimal Structural Guidance. Nano Lett. 17, 3782–3791 (2017).

62. Mezger, T. The Rheology Handbook. (Vincentz Network, 2020).

63. Janmey, P. A., Georges, P. C. & Hvidt, S. Basic Rheology for Biologists. in Cell Mechanics vol. 83 1– 27 (Elsevier, 2007).

64. Magnetic Resonance Imaging (MRI) of the Spine | HealthLink BC. https://www.healthlinkbc.ca/healthwise/magnetic-resonance-imaging-mri-spine.

65. Hwooi, S. K. W. & Sabri, A. Q. M. Enhanced correlation coefficient as a refinement of image registration. in 2017 IEEE International Conference on Signal and Image Processing Applications (ICSIPA) 216–221 (IEEE, 2017). doi:10.1109/ICSIPA.2017.8120609.

66. Junior, O. L., Delgado, D., Goncalves, V. & Nunes, U. Trainable classifier-fusion schemes: An application to pedestrian detection. in 2009 12th International IEEE Conference on Intelligent Transportation Systems 1–6 (IEEE, 2009). doi:10.1109/ITSC.2009.5309700.

67. Mohapatra, S. & Weisshaar, J. C. Modified Pearson correlation coefficient for two-color imaging in spherocylindrical cells. BMC Bioinformatics 19, 428 (2018).

68. Napoli, N. J., Barnes, L. E. & Premaratne, K. Correlation coefficient based template matching: Accounting for uncertainty in selecting the winner. in 311–318 (IEEE, 2015).

69. Kim, J. & Fessler, J. A. Intensity-based image registration using robust correlation coefficients. IEEE Trans. Med. Imaging 23, 1430–1444 (2004).

70. Brown, L. G. A survey of image registration techniques. ACM Comput. Surv. 24, 325–376 (1992).

71. Asuero, A. G., Sayago, A. & González, A. G. The correlation coefficient: an overview. Critical Reviews in Analytical Chemistry 36, 41–59 (2006).

72. Cameron, T. C. et al. PDMS Organ-On-Chip Design and Fabrication: Strategies for Improving Fluidic Integration and Chip Robustness of Rapidly Prototyped Microfluidic In Vitro Models. Micromachines (Basel) 13, (2022).

73. Wang, Z., Bovik, A. C., Sheikh, H. R. & Simoncelli, E. P. Image quality assessment: from error visibility to structural similarity. IEEE Trans. Image Process. 13, 600–612 (2004).

74. Sara, U., Akter, M. & Uddin, M. S. Image Quality Assessment through FSIM, SSIM, MSE and PSNR— A Comparative Study. JCC 07, 8–18 (2019).

75. Ciric, D. G., Peric, Z. H., Milenkovic, M. & Vucic, N. J. Evaluating Similarity of Spectrogram-like Images of DC Motor Sounds by Pearson Correlation Coefficient. ELEKTRON. ELEKTROTECH. 28, 37–44 (2022).

76. Blaney, L. Magnetite (Fe3O4): Properties, Synthesis, and Applications. (2007).

77. Oleic acid, 112-80-1, O1008, Sigma-Aldrich. https://www.sigmaaldrich.com/CA/en/product/sial/o1008.

78. Catauro, M., Bollino, F., Cristina Mozzati, M., Ferrara, C. & Mustarelli, P. Structure and magnetic properties of SiO2/PCL novel sol–gel organic–inorganic hybrid materials. J. Solid State Chem. 203, 92– 99 (2013).

79. Polycaprolactone average Mn 80,000 2-Oxepanone homopolymer. https://www.sigmaaldrich.com/CA/en/product/aldrich/440744.

80. Mohr, P. J., Newell, D. B. & Taylor, B. N. CODATA recommended values of the fundamental physical constants: 2014. Rev. Mod. Phys. 88, 035009 (2016).

81. Mulay, L. N. Basic concepts related to magnetic fields and magnetic susceptibility. in Biological effects of magnetic fields (ed. Barnothy, M. F.) 33–55 (Springer US, 1995). doi:10.1007/978-1-4757-0214-9_3.

82. Jones, R. M. Micromechanical Analysis of a Lamina. in Mechanics of Composite Materials (1998).

83. Tam, D. K. Y., Ruan, S., Gao, P. & Yu, T. High-performance ballistic protection using polymer nanocomposites. in Advances in military textiles and personal equipment 213–237 (Elsevier, 2012). doi:10.1533/9780857095572.2.213.

84. Zheng, J., Chen, S. & Xiao, Y. Effective permeability model of magnetorheological fluids and its experimental verification. J. Magn. Magn. Mater. 562, 169774 (2022).

