## Supplementary Information for "Effects of MRI on an Injectable Hydrogel with Magnetically Alignable Microstructures for Oriented Cell Growth"

**Supplementary Table S1.** List of variables used in this work.

| Variable | Definition | Unit |
|---|---|---|
| $\mathbf{M}$ | Magnetization Vector | A/m; emu/g |
| $\mathbf{M}_s$ | Saturation Magnetization Vector | A/m; emu/g |
| $M_s$ | Saturation Magnetization (Magnitude) | A/m; emu/g |
| $\chi$ | Magnetic Susceptibility | - |
| $\mathbf{H}$ | Internal Magnetic Field Vector | |
| $\mathbf{H}^{sat}$ | Saturation Internal Magnetic Field Vector | A/m |
| $\tau_{mag}$ | Magnetic Torque | N m |
| V | Volume | $m^3$ |
| $B_0$ | Applied Magnetic Field; in this work this is the applied MRI DC Field | T |
| $D_a$ | Demagnetizing Factor, along the axis of symmetry | - |
| $D_r$ | Demagnetizing Factor, radial | - |
| $\mu_0$ | Permeability of Free Space | H/m; N/A |
| $\beta$ | Angle between Applied Magnetic Field and the Direction of Magnetization; in this case, along the rod axis of symmetry | Radians (rads) |
| $\beta_0$ | Initial $\beta$ | Rads |
| $\phi_{SPION}$ | Volume Percent of SPIONs in Rods | % |
| $\rho_{SPION}$ | Density of SPIONs | $g/cm^3$ |
| $\tau_{elastic}$ | Elastic Torque | N m |
| G’ | Elastic Shear (storage) Modulus | Pa |
| G’’ | Loss Modulus | Pa |
| $\omega_r$ | Angular Frequency for Rheological Tests | Rad/s |
| L | Rod Length | m |
| D | Rod Diameter | m |
| p | Anisotropy Ratio, L/D | - |
| g(p) | Dimensionless Function of the Anisotropy Ratio, p | - |
| $\theta$ | Rod Angle of Deflection with Respect to its Initial Angle, $\beta_0$ ; this initial angle is relative to the applied magnetic field direction | Rads |
| $\tau_{viscous}$ | Viscous Torque | N m |
| $\eta_0$ | Zero- shear Viscosity of Gel | Pa s |
| $\tau_{inertia}$ | Inertial Torque | N m |
| I | Moment of Inertia | $kg\ m^2$ |
| P | Power Generated in a Spherical Sample (in this work, due to RF field induced eddy current) | W |
| R | Radius | m |
| $\sigma$ | Electrical Conductivity | S/m |
| D | Duty Cycle of RF Pulse | % |
| $\omega$ | Angular Frequency of the RF Magnetic Field | Rad/s |
| $B_1$ | RF Magnetic Field Amplitude | T |
| $\Delta T$ | Temperature Change | °C |
| Q | Heat Absorbed by Sample | J |
| m | Mass | kg |
| c | Specific Heat Capacity | J/ (kg K) |
| $B_{eff}$ | Effective Rectangular Pulse Amplitude | T |
| PW | Pulse Width | s; ms |
| TR | Repetition Time | s; ms |
| $\theta_f$ | Flip Angle | Rads |
| $\gamma$ | Gyromagnetic Ratio | Hz/T |

### I. PDMS Device Fabrication

#### a. 4-well chip

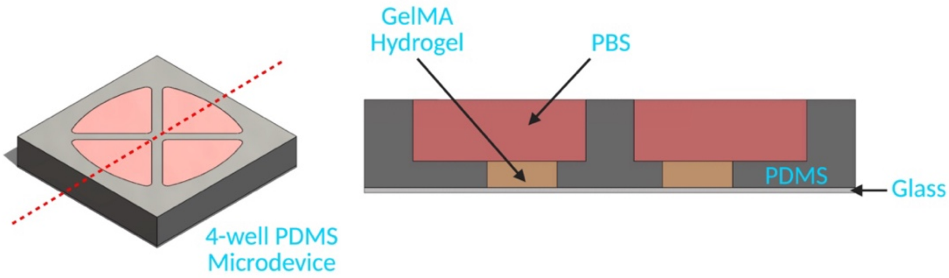


**Supplementary Figure S1.** Layers of 4-well microdevice. The microdevice consists of a Polydimethylsiloxane (PDMS) gasket with patterned well and reservoir features bonded to a glass coverslip. Each microdevice contains 4 independent hydrogel wells so that liquid GelMA can be loaded independently into each well but then simultaneously exposed to the same light source and magnetic field, sequentially. Overlying reservoirs can be filled with PBS to maintain hydration of the hydrogel during experimental scans. (Created in BioRender. Bennet, T. (2026) https://BioRender.com/agnd3h0)

Polydimethylsiloxane (PDMS)-glass microdevices containing 4 independent microwells were fabricated through replica molding and assembled via oxygen plasma treatment. Each well contains a small central hydrogel reservoir (4 mm diameter, 1.65 mm height) with a larger media reservoir directly on top as shown in Fig. S1. To create the PDMS layers containing the well features, liquid PDMS (Sylgard 184, Dow Corning) mixed at a 10:1 (elastomer base: curing agent) ratio was poured into molds created using 3D printing (MiiCraft). A transparency film lid was placed over the PDMS-filled molds after degassing prior to curing to obtain a uniform thickness as previously described in [72]. PDMS layers were cured overnight at 65°C.

#### b. Mock Spinal Lesion (MSL)

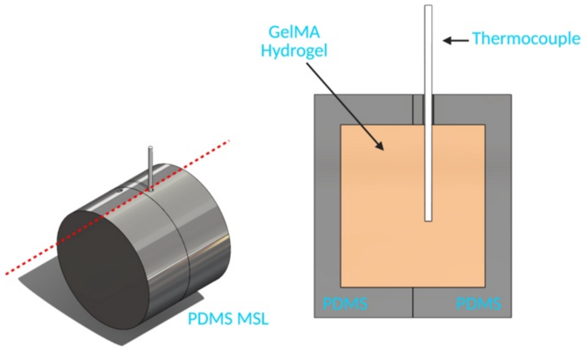


**Supplementary Figure S2.** Layers of Mock Spinal Cord Lesion (MSL). PDMS surrounds a GelMA filled cavity and contains an access point for the thermocouple. (Created in BioRender. Bennet, T. (2026) https://BioRender.com/085kpdt)

As shown in Fig. S2, MSLs were fabricated from PDMS using replica molding. Due to the limitations of the replica molding process, MSLs were fabricated by bonding two PDMS layers with well-like features together to create an enclosed 1 mL empty lesion cavity that can be filled with the desired hydrogel. To facilitate insertion of a thermocouple for thermal measurements, a 20G dispensing tip (Outer diameter: 0.9 mm; Cat.75165A677, McMaster Carr) was inserted into a hole in the mold for the PDMS layer during the casting process. Once the mold was assembled and the needle inserted, liquid PDMS (10:1 ratio) was poured into the resin molds and degassed for 30-40 min in a vacuum chamber prior to curing. A transparency film lid was used as previously described to obtain uniform thickness and to facilitate assembly[72]. Once cured, the PDMS MSL halves were removed from the molds and bonded together. To prevent leakage, the assembled MSLs were enrobed in liquid PDMS. Care was taken to ensure no liquid PDMS entered the empty cavity through the thermocouple access point.

### II. Gel Characterization Rheological Tests

(a)

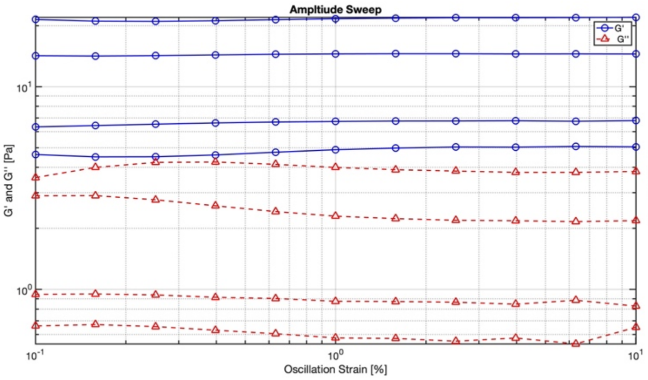


(b)

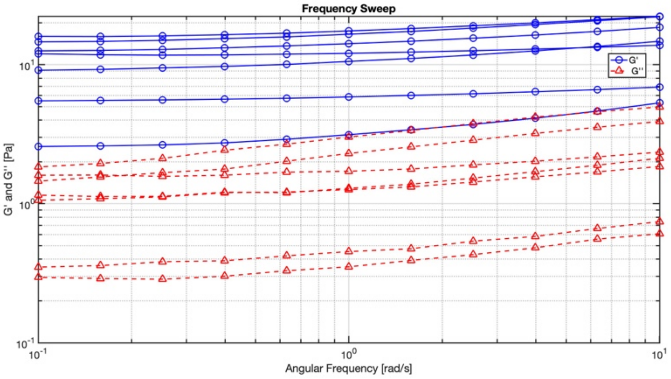


(c)

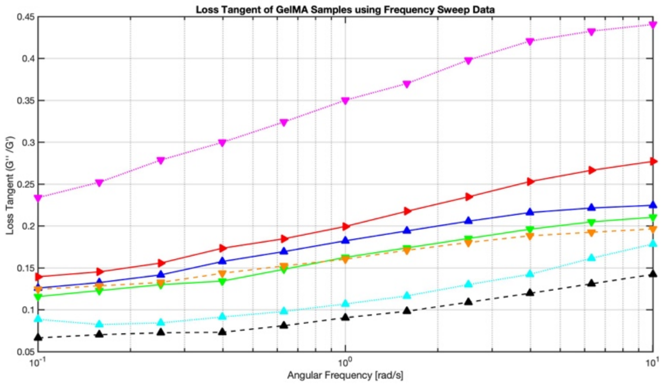


(d)

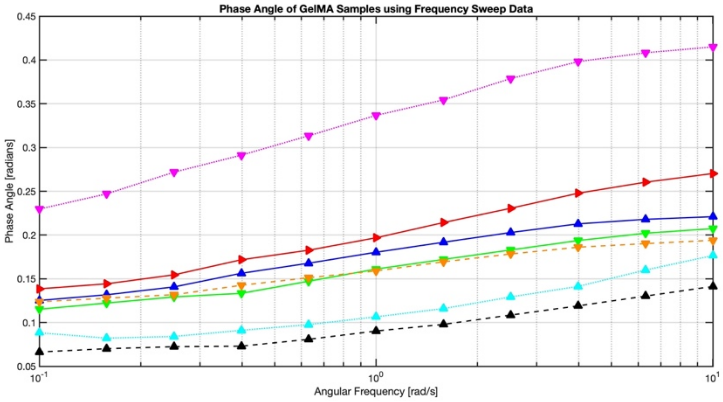


**Supplementary Figure S3.** Rheological test results of 6% GelMA, 50% DoF. **(a)** Amplitude sweeps data (n = 4); **(b)** frequency sweep data (n = 7); **(c)** loss tangent using frequency sweep data (n = 7), **(d)** phase angle using frequency sweep data (n = 7).

### III. Alignment State Study: Numerical Quantification Methods

**Supplementary Table S2.** Summary of numerical quantification methods used for pre- and post- MRI scan image comparison.

| Method | Range | Highest Similarity Value | Comparison Metric | Mathematical Algorithm | Rationale for Use |
|---|---|---|---|---|---|
| **RMSE** | 0 – 1 | 0 | Pixel Intensity | Square root of average (difference in pixel intensity between the two images)$^2$ | Widely used metric for comparison, based on the "error sensitivity philosophy" [73] comparing the images pixel by pixel [74]; limitations as discussed in [73] |
| **CC** | -1 – 1 | 1 | Pixel Intensity | Correlation Coefficient | Commonly used tool for image registration [69] and template matching [68] measuring the "degree of linear correlation in pixel-by-pixel intensity" [67]; additional advantages and limitations outlined in [75] |

### IV. Alignment State Study: Results

**Supplementary Table S3**. Numerical comparison values of chips 1-8 which underwent an MRI scan and the control chip (chip 9). This data is presented as the mean, with SD referring to standard deviation. The median can be found in Fig. 2(c).

| Chip | Chip Condition | RMSE Average ± SD | CC ± SD |
|---|---|---|---|
| **1, 5** | Unaligned, ‖ | 0.130 ±0.056 | 0.69 ±0.21 |
| **2, 6** | Aligned, ‖ | 0.15 ±0.052 | 0.65 ±0.22 |
| **3, 7** | Unaligned, ⊥ | 0.13 ±0.044 | 0.70 ±0.20 |
| **4, 8** | Aligned, ⊥ | 0.15 ±0.053 | 0.57 ±0.25 |
| **9** | Aligned, no MRI (Control) | 0.16 ±0.050 | 0.58 ±0.23 |

### V. Heat Study: Results

Results from the three 45-minute MRI scans conducted can be seen in Fig. S3. The averaged results can be found in Fig. 3(b) in the main text.

**(a)**

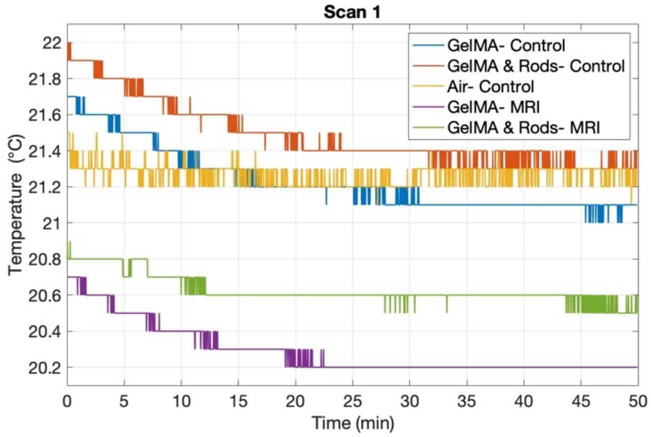


**(b)**

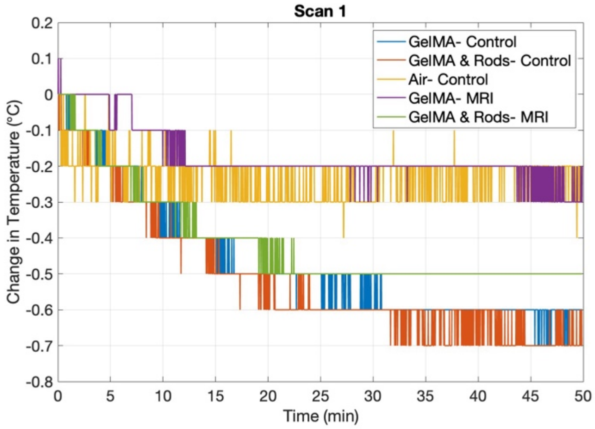


**(c)**

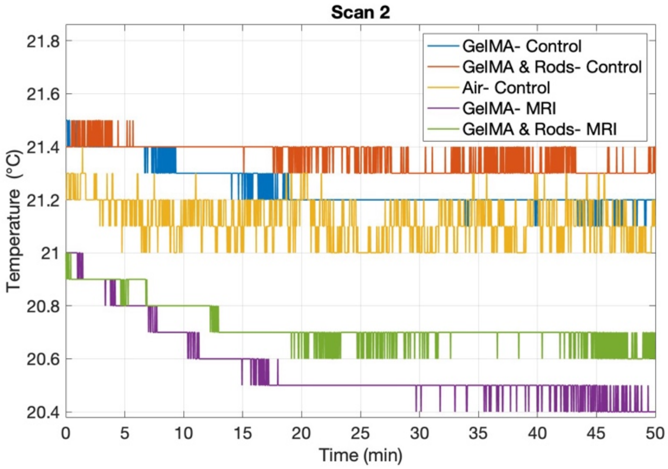


**(d)**

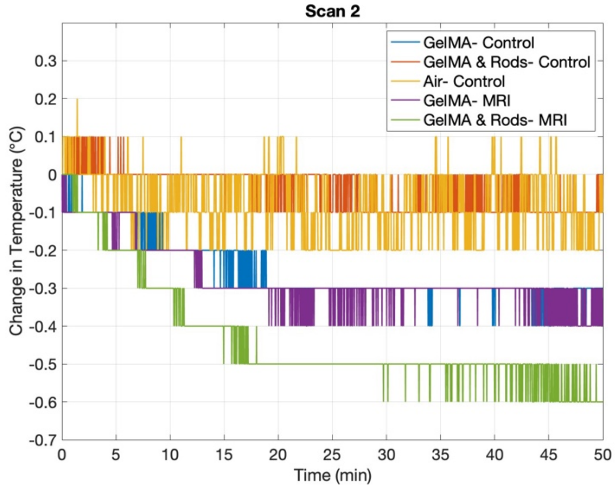

(e) (f)

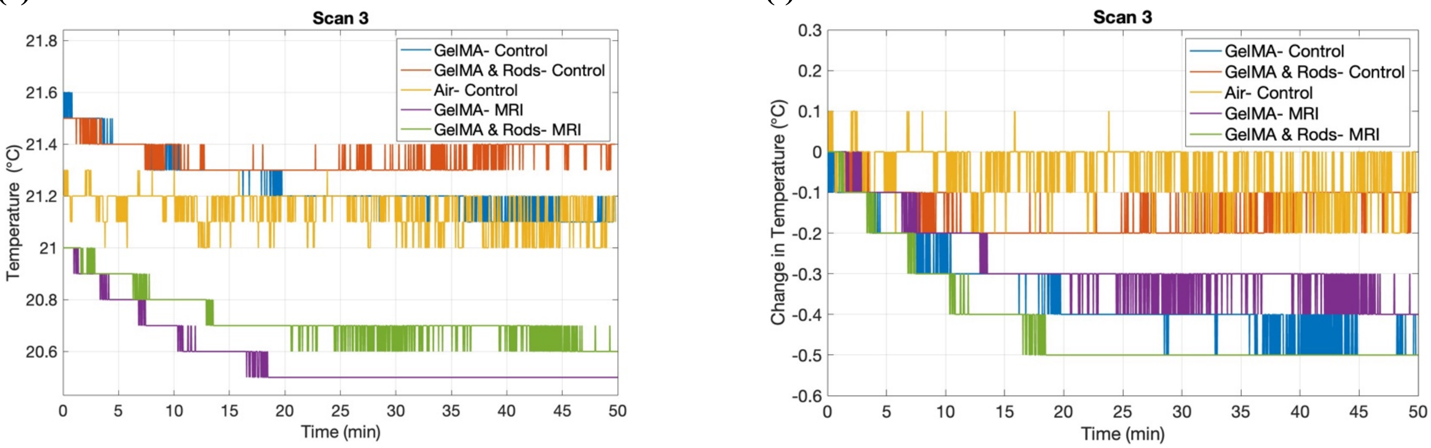


**Supplementary Figure S4.** Temperature and change in temperature of the samples over the course of a 45-minute MRI scan. The time spans to 50 minutes as there are pauses in between scans leading to the total scan time being 50-minutes. The sampling rate of the measurements is 1 second. The change in temperature values is relative to the starting temperature of each sample respectively. **(a)** Temperature of samples in the first scan. **(b)** Change in the temperature of the samples in the first scan. **(c)** Temperature of samples in the second scan. **(d)** Change in the temperature of the samples in the second scan. **(e)** Temperature of samples in the third scan. **(f)** Change in the temperature of the samples in the third scan.

### VI. Additional Rod Constants Calculations: Effective Magnetic Susceptibility

SPION Density ($\rho_{SPION}$) and Volume Fraction ($\phi_{SPION}$) in PCL Rods:

Given the below constants:

- SPION weight percent of final rod wight, $w_{SPION}$ = 0.99 weight(w/w) %
- SPION properties:
  - Initial volume magnetic susceptibility, $\chi_{SPION}^{vol}$ = 0.2 (EMG 1200, Ferrotec)
  - Components (EMG 1200, Ferrotec):
    - Magnetite, $Fe_3O_4$:
      - $w_{Fe_3O_4}$ = 60 - 80 w/w%
      - density, $\rho_{Fe_3O_4}$ = 5.18 g/cm$^3$ [76]
    - Fatty acid coating (oleic acid):
      - $w_{OA}$ = 20 - 40 w/w%
      - $\rho_{OA}$ = 0.89 g/cm$^3$ [77]
- PCL properties:
  - mass susceptibility, $\chi_{PCL}^{mass}$ = -7.5 × 10$^{-9}$ m$^3$/kg [78]
  - $\rho_{PCL}$ = 1.145 g/cm$^3$ [79]
- Additional relevant constant:
  - vacuum permeability, $\mu_0 = 4\pi \times 10^{-7}$ H/m [80]

The volume magnetic susceptibility of the PCL is calculated as outlined in equation (S1) below [81]:

$$\chi_{PCL}^{vol} = \chi_{PCL}^{mass}\rho_{PCL} = (-7.5 \times 10^{-9}\ \text{m}^3/\text{kg})\ (1.145\ \text{g/cm}^3) * 10^{-3}\ \text{kg/g} * 10^{6}\ \text{cm}^3/\text{m}^3 = -8.59\ \text{ppm}, \quad \text{(S1)}$$

From the SPION properties listed above, the effective SPION density can be calculated using the rule of mixtures, equation (S2) [82,83]:

$$\rho_{composite} = \sum \phi_i \rho_i = \frac{1}{\sum \frac{w_i}{\rho_i}} \quad \text{, so } \rho_{SPION} = \frac{1}{\frac{w_{Fe_3O_4}}{\rho_{Fe_3O_4}} + \frac{1-w_{Fe_3O_4}}{\rho_{OA}}}, \quad \text{(S2)}$$

For $w_{Fe_3O_4} = 0.60$, $\rho_{SPION} = 1.77$ g/cm$^3$, while for $w_{Fe_3O_4} = 0.80$, $\rho_{SPION} = 2.64$ g/cm$^3$.

For SPION density, the average value is used for calculations: $\rho_{SPION}$ ~2.21 g/cm$^3$.
Similarly, the density of a microrod can be calculated as shown in equation (S3):

$$\rho_{rod} = \frac{1}{\frac{w_{\text{SPION}}}{\rho_{\text{SPION}}} + \frac{1-w_{\text{SPION}}}{\rho_{PCL}}}, \qquad \text{(S3)}$$

Converting the weight% of SPIONs to volume% (vol%, $\phi$) [82]:

$$\phi_{SPION} = \frac{w_{SPION}}{\rho_{SPION}} \rho_{rod} = \frac{\frac{w_{SPION}}{\rho_{SPION}}}{\frac{w_{SPION}}{\rho_{SPION}} + \frac{1-w_{SPION}}{\rho_{PCL}}}, \qquad \text{(S4)}$$

$\phi_{SPION}$ ~ $0.0043 - 0.0064$ $(0.43 - 0.64$ vol%)
The average value is therefore used for the calculations: $\phi_{SPION}$ ~ $0.54\ vol\%$.

Rod Effective Magnetic Susceptibility:

The effective magnetic susceptibility of the rod can be determined by calculating the effective magnetic permeability of the SPIONs, PCL, and rod before converting the rod magnetic permeability to magnetic susceptibility. The effective magnetic permeability ($\mu_{eff}, ie.\ \mu_{rod}$) of a low concentration magnetic mixture with spherical magnetic particles can be calculated using the Maxwell-Garnett mixing model of magnetorheological fluids, as presented in equation (S5) [8481]:

$$\mu_{rod} = \mu_{PCL} + \frac{3\phi_{SPION}\, \mu_{PCL}\, (\mu_{SPION} - \mu_{PCL})}{\mu_{SPION} + 2\mu_{PCL} - \phi_{SPION}\ (\mu_{SPION} - \mu_{PCL})}, \qquad \text{(S5)}$$

where $\mu_i = \mu_0(1 + \chi_i^{vol})$ with *i* being PCL, SPION, and rod.
Given the range in volume fraction of SPIONs, the above equation leads to $\mu_{rod} = 1.258 \times 10^{-6}$ H/m and $\chi_{rod}^{vol} = 0.00080 - 0.0012$.
Alternatively, using the volume% of SPIONs and PCL in the rods, along with the volume magnetic susceptibility of the SPIONs and PCL, the effective magnetic susceptibility of a single rod can be calculated using the rule of mixtures. The Maxwell-Garnett mixing model introduced earlier considers the demagnetization factor of the spherical SPIONs (1/3) and incorporates this into the equation. For a simple mixing model, the apparent magnetic susceptibility of the SPIONs considering the demagnetization factor needs to be calculated, as presented in equation (S6-7):

$$\chi_{SPION,APP} = \frac{\chi_{SPION}^{vol}}{1+\frac{1}{3}\chi_{SPION}^{vol}}, \qquad \text{(S6)}$$

$$\chi_{rod}^{vol} = \phi_{SPION}\, \chi_{SPION,\text{APP}}^{vol} + (1 - \phi_{SPION})\, \chi_{PCL}^{vol} = 0.00080 - 0.0012, \qquad \text{(S7)}$$

The two methods lead the same results: $\chi_{rod}^{vol} = 0.00080 - 0.0012$. For calculations, the average value of 0.0010 can be used.

Rod Demagnetizing Factors:

Approximating the rods as very slender prolate ellipsoids (in fact, prolate spheroids as two axes are equal), the demagnetizing factors of the rods, along the axis of symmetry and radially, can be calculated as $D_a$ = 0.0558, $D_r$ = 0.4721, respectively [53].

### VII. MRI Scan Variables

**Supplementary Table S4.** MRI scan variables used to determine the expected heating, as shown in Table S5. $\theta_f$: Flip Angle, PW: Pulse Width, TR: Repetition Time.

| | Excitation Pulse | | | Refocusing Pulse | | | Fat Suppression Pulse | | | TR (ms) |
|---|---|---|---|---|---|---|---|---|---|---|
| Scan Name | $\theta_f$ (°) | PW (ms) | Pulses per TR | $\theta_f$ (°) | PW (ms) | Pulses per TR | $\theta_f$ (°) | PW (ms) | Pulses per TR | |
| FLASH 1,2,3: Initial three-slice localizer | 10 | 1.1100 | 1 | - | - | - | - | - | - | 15.00 |
| RARE 1: T2_TurboRARE_ax | 90 | 2.1000 | 2 | 180 | 1.7000 | 9 | - | - | - | 2500.00 |
| RARE 2: T2_Turbo_RARE | 90 | 2.1000 | 2 | 180 | 1.7000 | 9 | - | - | - | 2500.00 |
| RARE 3: T2_Turbo_RARE | 90 | 2.1000 | 2 | 180 | 1.7000 | 9 | - | - | - | 2500.00 |
| FLASH 1: T1_FLASH_3D_iso _quick | 20 | 3.5000 | 1 | - | - | - | - | - | - | 50.00 |
| FLASH 2: T1_FLASH_3D_iso | 20 | 3.5000 | 1 | - | - | - | - | - | - | 50.00 |
| EPI 1: SE_EPI | 90 | 1.9091 | 1 | 180 | 1.5455 | 1 | 90 | 1.9571 | 1 | 1500.00 |
| EPI 2: SE_EPI | 90 | 1.9091 | 1 | 180 | 1.5455 | 1 | 90 | 1.9571 | 1 | 1500.00 |
| DT-EPI: DTI_EPI_3D_40dir | 90 | 2.7273 | 1 | - | - | - | 90 | 1.9571 | 1 | 2100.00 |
| UTE 1: UTE2D | 15 | 0.3938 | 1 | - | - | - | - | - | - | 30.00 |
| UTE 2: UTE3D | 5 | 0.0043 | 1 | - | - | - | - | - | - | 4.24 |

**Supplementary Table S5**. Expected temperature rise by scan type. Explanation of the scans can be found in the *Methods* (Section 4.4.1).

| Scan Name | Total Average RF Power, P ($\mu W$) | Total Time of Scan (s) | Heat, Q (mJ) | ΔT (°C) |
|---|---|---|---|---|
| FLASH 1,2,3: Initial three-slice localizer | 0.328 | 43.00 | 0.0141 | $3.37 \times 10^{-6}$ |
| RARE 1: T2_TurboRARE_ax | 3.92 | 80.00 | 0.313 | $7.49 \times 10^{-5}$ |
| RARE 2: T2_Turbo_RARE | 3.92 | 80.00 | 0.313 | $7.49 \times 10^{-5}$ |
| RARE 3: T2_Turbo_RARE | 3.92 | 40.00 | 0.157 | $3.74 \times 10^{-5}$ |
| FLASH 1: T1_FLASH_3D_iso_quick | 0.125 | 89.00 | 0.0111 | $2.66 \times 10^{-6}$ |
| FLASH 2: T1_FLASH_3D_iso | 0.125 | 2.00 | 0.000250 | $5.97 \times 10^{-8}$ |
| EPI 1: SE_EPI | 1.07 | 2.00 | 0.00214 | $5.11 \times 10^{-7}$ |
| EPI 2: SE_EPI | 1.07 | 450.00 | 0.481 | $1.15 \times 10^{-4}$ |
| DT-EPI: DTI_EPI_3D_40dir | 0.185 | 840.00 | 0.155 | $3.71 \times 10^{-5}$ |
| UTE 1: UTE2D | 1.04 | 13.00 | 0.0135 | $3.23 \times 10^{-6}$ |
| UTE 2: UTE3D | 75.0 | 55.00 | 4.12 | $9.86 \times 10^{-4}$ |
